# A panoramic perspective on human phosphosites

**DOI:** 10.1101/2022.03.08.483252

**Authors:** Pathmanaban Ramasamy, Elien Vandermarliere, Wim vranken, Lennart Martens

## Abstract

Protein phosphorylation is the most common post-translational reversible modification of proteins and is key in the regulation of many cellular processes. Due to this importance, phosphorylation is extensively studied, resulting in the availability of a large amount of mass spectrometry based phospho-proteomics data. Here, we leverage the information in these large-scale phospho-proteomics datasets, as contained in Scop3P, to analyze and characterize proteome-wide protein phosphorylation sites (P-sites). First, we set out to differentiate correctly observed P-sites from false positive sites using five complementary site properties. We then describe the context of these P-sites in terms of protein structure, solvent accessibility, structural transitions and disorder, and biophysical properties. We also investigate the relative prevalence of disease-linked mutations on and around P-sites. Moreover, we also assess structural dynamics of P-sites in their phosphorylated and unphosphorylated state. Our study shows that the residues that gets phosphorylated are more flexible than their equivalent non-phosphorylated residues. Our structural and biophysical analyses of P-sites in solvent inaccessible (buried) regions of proteins show that these sites are primarily found in multi-site phospho-proteins, where highly dynamic structural transitions can occur upon binding with another protein. Finally, our analysis of the biophysical properties of P-site mutations shows that P-site mutations that occur in structurally rigid regions are more often involved in disease.

## Introduction

Proteins carry out most functions in the cell, and are carefully regulated to ensure correct operation of the cell. Correspondingly, dysregulation of protein function often leads to diseases, including cancer^1,2^. A common mechanism for protein regulation is through post-translational modifications (PTMs), which can affect the activity, sub-cellular localization, and stability of proteins. While many PTMs play important roles in cellular regulation, phosphorylation is one of the most prevalent and well-studied^3,4^. Phosphorylation of proteins can, amongst others, alter their activity, change their binding affinity for substrates, interfere with their interactions (protein-protein interactions), alter their conformation and localization, and control their degradation^5,6^. Phospho groups are typically transferred to the hydroxylated side chains of Ser, Thr or Tyr by kinases. Previously, it has been shown that at least 30% of all proteins contain at least one phosphorylated residue^7^. Ser is the most commonly phosphorylated residue, followed by Thr, while Tyr accounts for few phosphorylation events^8^. This is due to the high number of protein kinases that are specific to Ser and Thr residues (Ser/Thr kinases; 80% of known kinases^7^). Moreover, Tyr kinases are activated only under specific circumstances, unlike the more broadly active Ser/Thr kinases^8,9^.

Many proteins actually have multiple phosphorylation sites (P-sites), which play an important role by expanding the regulation by modulating conformational change favoring increased binding affinity to proteins^10^. This multi-site phosphorylation can occur through progressive phosphorylation of sites after initial binding of the kinase, or through non-progressive phosphorylation involving repeated binding and unbinding of the kinase^11^. In both cases, the order of site phosphorylation is based on the catalytic efficiency of the kinase^11,12^ kinase-substrate sequence and structural specificity^13^. In progressive phosphorylation, the distance between sequential P-sites is critical to maintain the phosphorylation process and the binding specificity of kinases, but exact distance constraints have not yet been determined^14^. On the other hand, non-progressive phosphorylation of a certain P-site can act as a signal to recruit the same (or even a different) kinase to the protein to phosphorylate a subsequent P-site in a process known as substrate “priming”^11,15,16^. Interestingly, highly efficient P-sites can also prevent the phosphorylation of low efficient sites because of competitive kinase binding^17^.

Indeed, kinases phosphorylate more than one substrate protein inside the cellular compartment, which typically compete. As a result, one substrate can act as a competitive inhibitor for the other^6,17^. Hence, an effective phosphorylation event depends on the relevant combination of kinase, phosphatase, and substrate abundance. While high-efficient substrate sites will only require a low kinase concentration to become phosphorylated, poorly efficient substrate sites will require a higher kinase concentration^17,18^. Moreover, kinase specificity is not always perfect, which can result in the off-target phosphorylation of sites in highly abundant proteins^19^.

Most P-sites have been shown to occur in unstructured regions of proteins and in regions outside a defined domain^20,21^, most likely as these sites are highly accessible and visible to kinases. But phosphorylation can also occur in ordered or defined structural regions^22^, where they are often related to conformational changes^23^ resulting in the exposure of previously inaccessible sites. Such conformational changes can even be triggered through phosphorylation-induced alterations in a protein’s conformational dynamics, with increased net negative charge of this protein through sequential phosphorylation leading to the conformational switch^24^. Moreover, it has also been shown that phosphorylation can shift conformational balance in regions prone to order/disorder transitions^25,26^. As a result, phosphorylation can alter the oligomeric state of a protein due to the conformational changes it induces^27,28^. Phosphorylation can also trigger conformational changes and/or protein binding events that result in this (previously accessible) P-site becoming buried^29^.

The most frequently used tool to study proteome-wide protein phosphorylation is mass spectrometry-based proteomics^30,31^. Most of these so-called phosphor-proteomics studies rely on phosphorylation enrichment strategies to identify as many phospho-peptides as possible, resulting in the identification of thousands of P-sites from a single experiment^31–33^. Despite its successes, there are issues with this approach, most notably the phosphorylation site localization problem^34,35^, which is the false identification of a peptide to be phosphorylated by mis-localizing a P-site in the peptide; an issue that is not helped by apparent phosphorylation ‘jumps’ in the instrument^36–38,^ where a phospho group can relocate during peptide fragmentation forming non-native phosphorylated peptides. In addition, establishing the stoichiometry of identified P-sites remains a challenge as well. Indeed, it remains very difficult to assess how many copies of a protein are phosphorylated at a given position at a given time^39^. Moreover, the functional importance of many of the identified P-sites is unclear, with many P-sites likely representing off-target sites on highly abundant substrates. Hence, it is necessary to carefully scrutinize these data to select the most reliable P-sites, which can then be integrated with existing knowledge and *in silico* predictions to understand their functional role.

Such an analysis is important, as, despite increasing numbers of identified P-sites and evidence that most human proteins are phosphorylated, only 5% of P-sites have a known role^40^. Several approaches have therefore been applied to functionally prioritize relevant P-sites based on their annotation in terms of functional domains^41,42^, structure^21,43^, mutational analysis^44^, and computational exploration^20,45^. Several resources have been built around human phosphorylations; Scop3P^46^ is a database of human P-sites that are obtained from both re-processed human proteomics projects (PRIDE^47^) and UniProtKB/Swiss-Prot^48^ annotations. UniProtKB/Swiss-Prot^48^ contains annotated P-sites at three possible levels of evidence: experimental evidence described in literature (*experimental*); similarity from sequence homologs (*similarity*); or a combination of both (*combined*). dbPTM^49^ is a resource based on the integration of P-sites obtained from different resources like UniProtKB/Swiss-Prot and PhosphoELM^50^ while the P-sites in PhosphositePlus (PSP)^51^ are obtained from P-sites identified through low throughput (LTP) and high throughput (HTP) experiments. However, few, if any, of these studies have brought all these perspectives together to study proteome-wide protein phosphorylation in human samples.

In this work we performed an in-depth computational investigation of human P-sites obtained from the large-scale re-processing of multiple phospho-proteomic projects, as made available through Scop3P^46^. First, we differentiate true P-sites from noise by combining protein abundance, peptide evidence (singly or multi phosphorylated peptides), localization probability, distance between adjacent P-sites, and the frequency of P-sites observed in different phospho proteomic projects. We also identify preferential occurrence of P-sites in certain protein domains and structural regions. We performed a panoramic 360° investigation of P-sites and phosphoproteins by combining different parameters such as sequence, secondary structure properties, 3D coordinates, and biophysical properties such as backbone and side chain dynamics, disorder propensity, and early folding properties. We also investigated sites within regions of possible structural transitions to see if this co-occurrence might explain the presence of P-sites in non-solvent exposed regions. Finally, we also investigate the impact of single amino acid variation on proteins specifically for P-sites compared to non-phosphorylated STY sites.

## Methods

All data analyses and plots were done in Python (version 3) with the ‘pandas’ package. Protein structure figures were generated using PyMol^52^ (version 2.3.0). All the data and python scripts used in these methods are available online (https://github.com/Pathmanaban/Phospho_analysis).

### P-site data collection, initial filtering, and P-site occupancy

We retrieved all annotated P-sites from Scop3P^46^, PhosphositePlus^51^ (PSP)(v6.5.9.1) and dbPTM^49^ (data downloaded on Sep 2020). All obtained P-sites were remapped with UniProtKB/Swiss-Prot^48^ human sequences (release-2020_01 of 22 April 2020, with 20365 canonical sequences) for their correct position within the protein sequence. P-sites that still mismatched after this were excluded from further analysis. After remapping, 106,975 P-sites were retained out of 108,123 original P-sites from Scop3P.

Furthermore, only P-sites from Scop3P that were identified using reprocessed PRIDE data were considered. These sites already pre-filtered in Scop3P for having a site localization probability of ≥ 0.5. This resulted in 92,453 retained P-sites, which matched to 13,437 proteins.

We calculated phosphorylation occupancy for each of these protein based on the ratio of the number of P-sites identified in that protein to the total number of STY sites in a protein (F1).

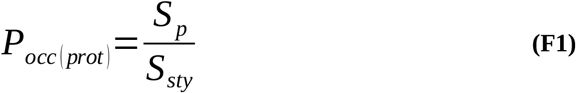

where S_p_ is the total number of experimentally identified P-sites in a protein, S_sty_ is the total number of STY sites in the same protein, representing all potential P-sites.

This resulted in 81,404 P-sites from 12,632 proteins, which were used as a **positive set**. Each of the sites in this positive sets are annotated with external resource evidence (UniProtKB/Swiss-Prot, dbPTM, PSP) for any overlap on phosphorylation annotation.

For the **negative set**, we retrieved all STY sites from the same 12,632 proteins used in the positive group for which evidence for phosphorylation is present in any of the four databases (Scop3P, UniProtKB/Swiss-Prot, PSP and dbPTM). Randomly selected subsets of these sites were then generated to match the respective population sizes in the positive set (*e.g.* number of S/T/Y P-sites), for use in different downstream analyses (sequence, biophysics and functional domains and structures, see the respective sections below for more details).

### Protein abundance calculation

Protein abundance was estimated using the normalized spectral abundance factor (NSAF), which allows comparison of individual proteins between different samples. NSAF is calculated using the formula F2.

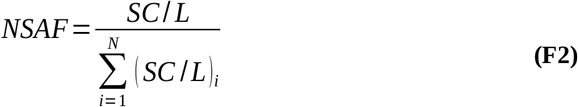

Where SC is the spectral count of the protein, *L* is the length of the protein and *N* is the total number of proteins identified in the experiment. Because these data are obtained across multiple experiments, an overall average *NSAF_avg_* value was calculated for each protein across all projects in which that protein was identified using the formula F3.

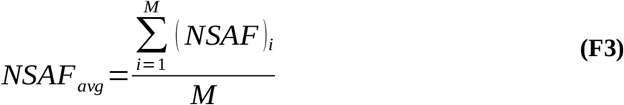

Where (NSAF)_i_ is the abundance of the protein in project_i_, M is the total number of projects a protein have been identified.

### Phosphorylation in functional domains

We retrieved domain annotations for human proteins from Pfam^53^ (release 33.1) and considered only Pfam-A entries, which contain curated information. P-sites were mapped onto Pfam domains scanning domain boundaries (Pfam start and Pfam end) for a particular protein.

Phosphorylation occupancy on domains was then calculated based on the number of P-sites in a domain compared to the total number of STY sites within this domain. However, because domain boundaries and domain sequence composition vary between proteins, the phosphorylation occupancy may differ for the same domain in different proteins. We therefore estimated the average phosphorylation occupancy as the ratio of the sum of all P-sites for that domain from across all proteins instances, to the sum of all STY sites in that domain across all protein instances (F4).

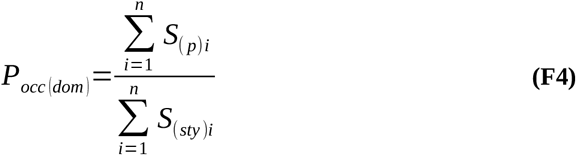

where **S**_(p)i_ and **S**_(sty)i_ are the total number of P-sites and total number of STY sites of the domain in a protein instance and **n** is the number of protein instances of a domain.

For example, for the actin domain (Pfam ID: PF00022), which is seen in 25 protein instances, the total sum of all P-sites across all 25 protein instances is 229, and the sum of all STY sites across all 25 protein instances is 1581. The average phosphorylation occupancy is then calculated as the ratio of total P-sites to total STY sites across all protein instances (229/1581)*100%, yielding 14.48%.

We also determined the overall phosphorylation contribution of a domain to the proteome, by the average number of sites phosphorylated in the domain compared to the total number of P-sites in our positive dataset (81,404 sites). For example, for the above mentioned actin domain, a total of 229 P-sites was obtained from all 25 proteins. The phosphorylation contribution of this particular domain to the proteome is then calculated as 229/81,404, or 0.28%.

### Disordered content and structural transition properties of phospho proteins and sites

Information on protein disorder, such as disordered sequence regions, regions involved in one or more structural states (order to disorder, disorder to order, molten globule, pre-molten globule), and phase transition regions were obtained from MobiDB^54^ (Version 4.0.1) and DisProt^55^ (release 2020_12). We only considered curated information from DisProt, and MobiDB-lite^56^ and IUPRED-long^57^ predictions from MobiDB. The union of the resulting two datasets was taken, with MobiDB contributing 53, 107 regions, and DisProt 169 regions.

P-sites were mapped onto these disordered regions and to simplify interpretation, we combined all ontology terms for a given P-site. For example, if a site is seen in a ‘disordered’ region as well as in a ‘disorder to order’ transition state region, we labeled the site with the combined term ‘disorder, disorder to order’ in the data.

### Biophysical properties and structural annotations

Biophysical property predictions were performed using DynaMine^58^ for backbone and side chain dynamics, DisoMine^59^ for disorder propensities, and EFoldMine^60^ for early folding properties. To understand the local environment of P-sites and non-P-sites, the same information was also obtained for the five residues before and after a site, totaling an eleven-residue fragment including the central amino acid.

### Structural properties

Protein structures and secondary structure properties were obtained from PDB^61^, through the Uniprot mappings already present in Scop3P. We only considered structures of at least 50 amino acids long, and furthermore only retained segments that mapped between UniProtKB/Swiss-Prot and PDB for at least 50 consecutive amino acids. To avoid redundancy, if a protein has more than one structure, the x-ray diffraction structure with the highest resolution is considered. If multiple structures share this highest resolution, the structure covering most of the P-sites is preferred. The structure coordinates were mapped to sequence using SIFTS^62^. Solvent accessibility and secondary structure information was obtained from PDBe-PISA^63^ and DSSP^64^, respectively. Secondary structural elements are classified into four categories (**H**: helices, **E**: beta strands, **C**: coils/unstructured, and –: no annotation/missing segments). This resulted in 8,432 P-sites from 3,059 phosphoproteins mapped onto 3,517 different structures.

The same data was obtained for the negative sets as described above. The negative sets were further filtered for the proteins in positive structure set such that it contained only non-P-sites from the same structures as the positive structure set (will be referred as **negative structure set**). These sites were sampled randomly to match the positive data points for statistical and downstream analysis. For both positive and negative sets the biological assembly assigned by PISA were used.

We classified residues as accessible (**A**), interface (**I**), or buried (**B**) based on the accessible surface area (ASA) and buried surface area (BSA) obtained from PDBe-PISA^63^ based on the following criteria:

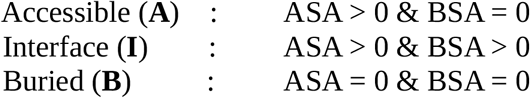

When analyzing P-sites in their structural context, biophysical properties like backbone dynamics, side chain dynamics, and early folding propensities were predicted on the respective PDB sequences.

To understand the local environment of P-sites and non-P-sites, the same information was also obtained for the five residues before and after a site, totaling an eleven-residue fragment including the central amino acid.

### Half-sphere exposure of residues

To obtain the residue contacts of the structural environment on and around P-sites and non-P-sites, we calculated the Half Sphere Exposure^65^ (HSE) of amino acids in structure. The HSE method calculates the number of spatially close residues by measuring residue contacts around a residue (usually <=13 Å) in the direction of its side chain (HSE up), as well as in the opposite direction towards the backbone (HSE down). Two residues are said to be in contact if their specific atoms (Cα or Cβ) are within a given radius (8 Å-14 Å)^66,67^. We calculated residue contacts for Cβ atoms around the P-site at a radius of 13Å^65,67^. Every site in the dataset has two values (HSE up and HSE down), each equal to the number of Cβ contacts for a given amino acid in that direction.

### Single amino acid variants (SAVs) in P-sites

We obtained single amino acid variation (SAVs) information for human proteins from the Humsavar information (Human polymorphisms and disease mutations, release 2020_06 of 02-Dec-2020) in UniProtKB/Swiss-Prot^48^. The obtained mutations were assigned two call labels (disease or neutral) based on their annotation in Humsavar. To understand the local environment of P-sites and non-P-sites, the same information was also obtained for the five residues before and after a site, totaling an eleven-residue fragment including the central amino acid. Mutations associated with P-sites were classified as ‘direct mutants’ (pSAVs) that fall on a P-site, ‘proximal’ if they are within two residues of the P-sites, or ‘distant’ if they are more than two residues away from the central P-site.

### Statistical significance calculations

To assess statistical significance for a given comparison between a positive (P-sites) and negative (non-P-sites) set, we make use of an empirical null distribution. This null distribution is created by randomly sampling a number of negative cases equal to the number of positive cases (negative cases always outnumber the positive cases), and then swapping the labels (positive and negative) randomly. The resulting distributions from the label-swapped data are then compared using the robust Kolmogorov–Smirnov test (KS test), yielding a p-value and a D-statistic for this label-swapped subset. This process of subsetting and label-swapping is then repeated 1000 times, yielding an empirical null distribution for both the *p*-value as well as the D-statistic, each consisting of 1000 values.

The KS test values from the comparison between the original positive and negative datasets are then compared to this empirical null distribution to test for significance. The significance is determined by the obtained D statistic and *p*-values by quantifying the distance between the empirical distribution of the two groups. *p*-values and D-statistics were determined using scipy (scipy. stats. ks_2samp).

## Results

We collected all P-sites from Scop3P which resulted in 106,975 P-sites in total, mapping to 14,145 human proteins in UniProtKB/Swiss-Prot. We compared the level of annotation of these Scop3P P-sites with these of two additional reference databases: from dbPTM and from PSP.

75% of the P-sites in Scop3P are also found in at least one of these two other databases. 65% of sites is found in all three databases (Scop3P, dbPTM and PSP), while 5% each are found in Scop3P & dbPTM, or in Scop3P & PSP **(Figure 1A i)**. Most of the sites that have combined evidence in UniProtKB/Swiss-Prot are also seen in re-processed projects (**Figure 1A ii).** We also observed that 65% of all P-sites from UniProtKB/Swiss-Prot were recovered through re-processing data in Scop3P and most of these P-sites have ‘combined’ evidence in UniProtKB/Swiss-Prot (**Figure 1A ii**).

**Figure 1:**
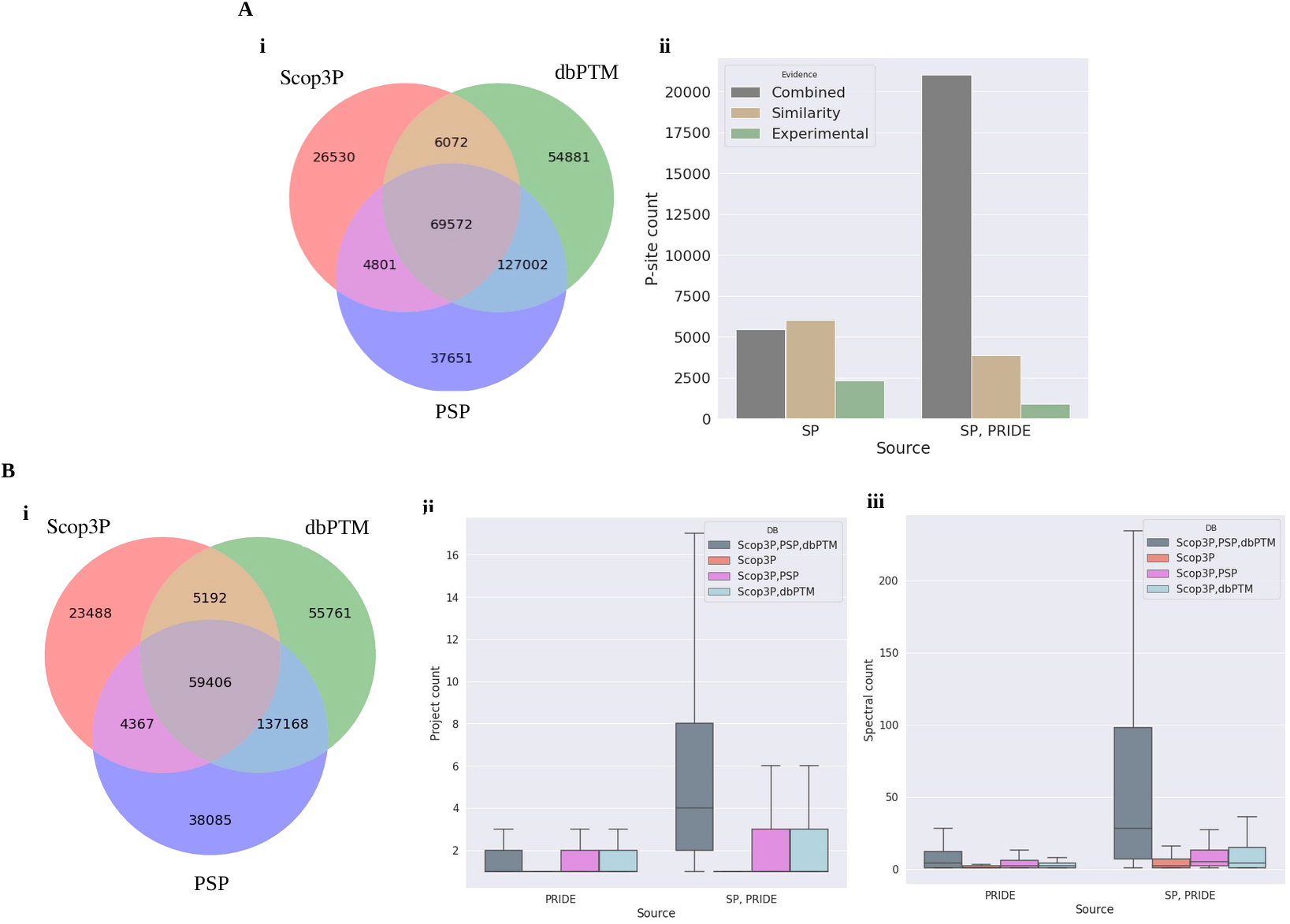
P-sites in Scop3P compared to other resources. **A) Scop3P P-sites from all resources** i) Total P-sites available from different resources and their common sites. ii) Annotations from UniProtKB showing the evidence of sites obtained by experimental methods, similarity from homologous sequences or combinatorial evidence (absolute counts shown in y-axis). ‘SP’ if the sites are from SwissProt and ‘SP, PRIDE’ if the sites are seen both in Swiss-Prot annotations and in re-processed PRIDE experiments. **B) Scop3P P-sites obtained only from re-processing proteomics projects** i) Total number of Re-processed P-sites that are common in three resources. ii) Frequency of P-sites with multiple project evidence and their database annotation iii) Frequency of number of spectra identified for the P-site from all re-processed experiments and their database annotation (y-axis shows the absolute count)

Next, to identify the reliable P-sites from random ones, we only considered P-sites identified by re-processed proteomics projects, as these sites have multiple metadata annotations like spectral evidence, protein abundance, site localization and P-site frequency across projects. Based on these restrictions, we continued our analysis with 92,453 P-sites from 13,437 human proteins that are identified by re-processing proteomics projects in Scop3P (**Figure 1Bi & Figure dataset overview**). When only re-processed sites were taken into account, we lose about 14,522 sites (annotations with evidence only from UniProtKB/Swiss-Prot). A similar concordance between the databases is found for this subset (**Figure 1B i**): 75% of the P-sites in Scop3P are also found in at least one of these two other databases (Scop3P, dbPTM and PSP; Scop3P & dbPTM; or Scop3P & PSP). We observed that P-sites that are identified in multiple proteomics projects are found more frequently in multiple databases (**Figure 1B ii**) and have higher spectral evidence. (**Figure 1B iii**).

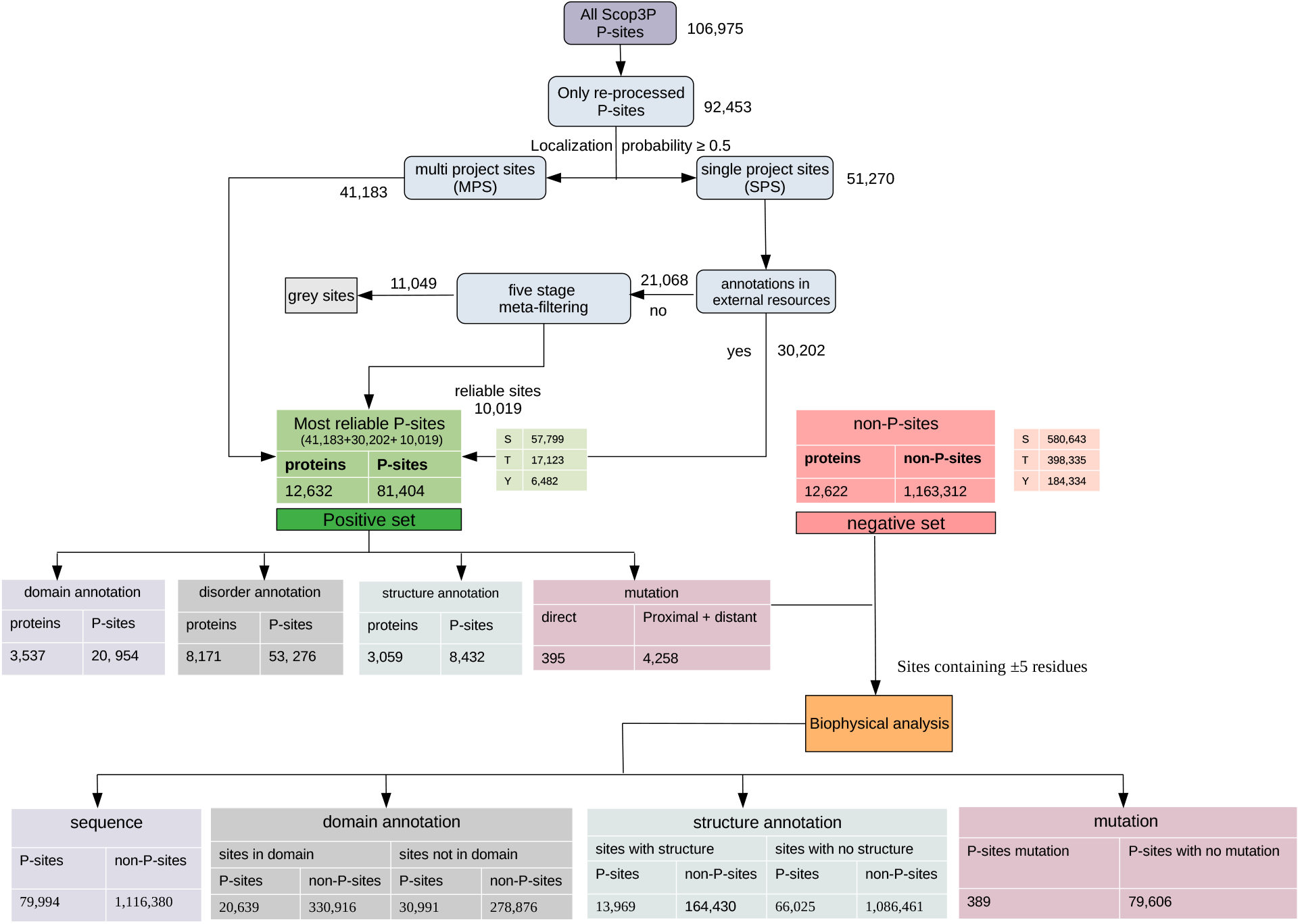
Figure dataset overview: Figure shows the overview of selecting positive and negative set for the analysis and the annotation of P-sites with domain, structure, mutation information used for different analysis in this work

Although phosphorylation is strictly regulated in multiple ways within the cell - cellular organization of kinases and phosphatases, and kinase substrate specificity - there still is a high chance that a random protein gets phosphorylated due to a random encounter of a kinase (off-target effects)^19^. We therefore compared the estimated abundance of proteins in relation to their phosphorylation occupancy. We found a moderate but positive linear relationship (Pearson correlation of 0.38) showing a significant (p<0.001) tendency for a larger number of identified P-sites with increasing protein abundance (**Figure 2A**) The correlation between protein abundance and phosphorylation status gives an indication of possible random phosphorylation events at protein level, but it does not provide any information on possible affected sites. Because these sites are identified from different phospho-proteomic experiments there might be a chance that a site from a given protein, identified in a given project **(Figure 2B)**, can be phosphorylated due to a random encounter, but the likelihood for a kinase to phosphorylate multiple copies of this site in that protein is taken to be low^19^, so the stoichiometry of these sites from the projects along with protein abundance should be a good measure to distinguish reliable P-sites from random ones. In order to see how many of these sites have occurred due to random effects we analyzed the sites in terms of the frequency of sites being seen as phosphorylated across different phospho-proteomic projects, the phosphorylation status of the peptide (singly/multiply phosphorylated), the phospho distance (distance between two identified P-sites), and the phospho localization probability.

**Figure 2:**
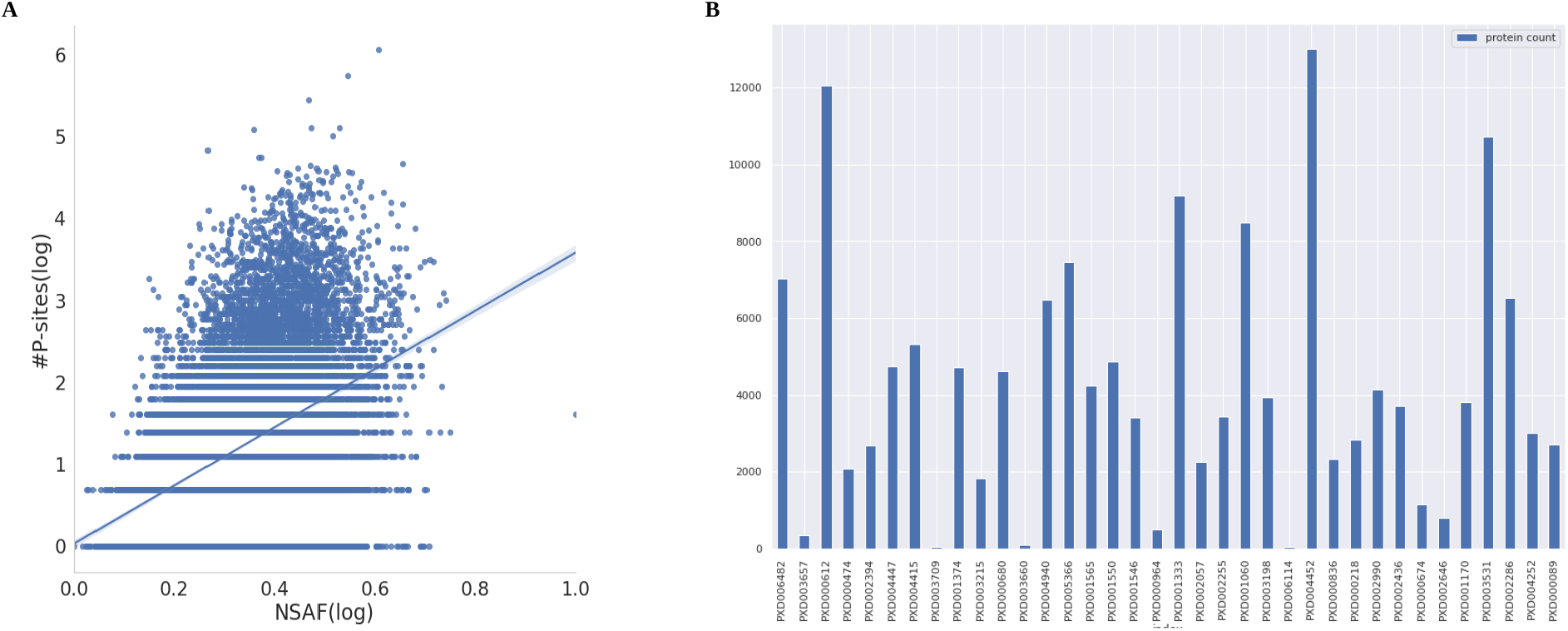
Phospho proteins identified through re-processing PRIDE projects. **A)** Protein abundance (NSAF) and the phosphorylation status of proteins. NSAF values of proteins (x-axis, log transformed) and the number of sites phosphorylated in the protein (y axis, log transformed). **B)** The total number of proteins identified per projects are shown. There are 36 different projects re-processed in Scop3P and the number of proteins identified differs based on the experimental goal of the original data submitted, from specific group of proteins (PXD000218), or enrichment strategies (PXD001060) or full proteome wide analysis (PXD000612). PRIDE project IDs in x-axis and number of protein identified per projects in y-axis (absolute count).

Our investigation on across-project frequency of P-sites showed that 55% of the sites (51,270 P-sites) are found to be phosphorylated in only one project (single project sites: SPS), while 45% of the P-sites (41,183 sites) are seen as phosphorylated in more than one project (multi project sites: MPS) (**Figure dataset overview**). A site is less likely to be identified as phosphorylated due to random chance in more than one phospho-proteomics experiments with higher localization probability (>0.7). So, we continued with the 51,270 SPS and try to filter them in different steps. As these sites are identified only in one project in our data, with varying localization probability and protein abundance, there is a high chance that these SPS might be random sites. We looked for possible bias for certain projects with more protein identifications (are these proteins and sites specific to certain projects) and found that the SPS and proteins containing SPS are not just coming from a particular project or experimental studies but are distributed across different projects (**Supplementary Figure S1**).

To further identify likely non-random SPS, we looked at how many of these have multiple database evidence. We found that 58% of the SPS (30,202 sites out of 51,270) are seen in Scop3P and at least one other database (23,327 or 45% in Scop3P, dbPTM and PSP; 3,805 or 7% in Scop3P & dbPTM; and 3,070 or 6% in Scop3P & PSP) (**Figure dataset overview**). Because these sites also have evidence of phosphorylation from at least one other resource along with re-processed evidence in Scop3P, we consider these to be reliable P-sites as well. We subsequently had a closer look using a five-stage filtering at the remaining 21,068 SPS that were very specific to Scop3P.

### Identifying reliable P-sites by applying five stage meta-filtering

We annotated the 21,068 SPS based on their protein abundance, stoichiometry of the P-site (frequency of phospho spectra identified for the particular site), localization probability, phosphorylation status, and phospho distance (the distance to the previous or next P-site on the same protein in primary amino acid sequence space). Labels were assigned based on the combination of the following criteria,

Based on the label combination we filtered the reliable P-sites; for example, sites that have label **S1D1L1P3A3 (Table 1)** are considered as reliable P-sites, while sites with label **S1D1L1P3A1 (Table 1)** are considered non-reliable. Both these groups are singly phosphorylated (**S1**), the nearest identified P-site is >20 amino acids away (**D1**), come with a localization probability of >70% (**L1**), and have <5 spectral frequency (**P3**). The only difference here is their protein abundance, as sites with label **S1D1L1P3A1** are from highly abundant proteins (**A1**) with fewer phospho spectra (**P3**), which can indicate phosphorylation due to random encounters with a kinase. Based on these combinations we conclude that 10,019 SPS (**Supplementary Figure S2: green bars and Table 1**) out of 21,068 Scop3P only sites are reliable P-sites. The remaining 11,049 sites were classified to be in the “grey zone”; as these sites are not easily substantiated.

**Table 1:**
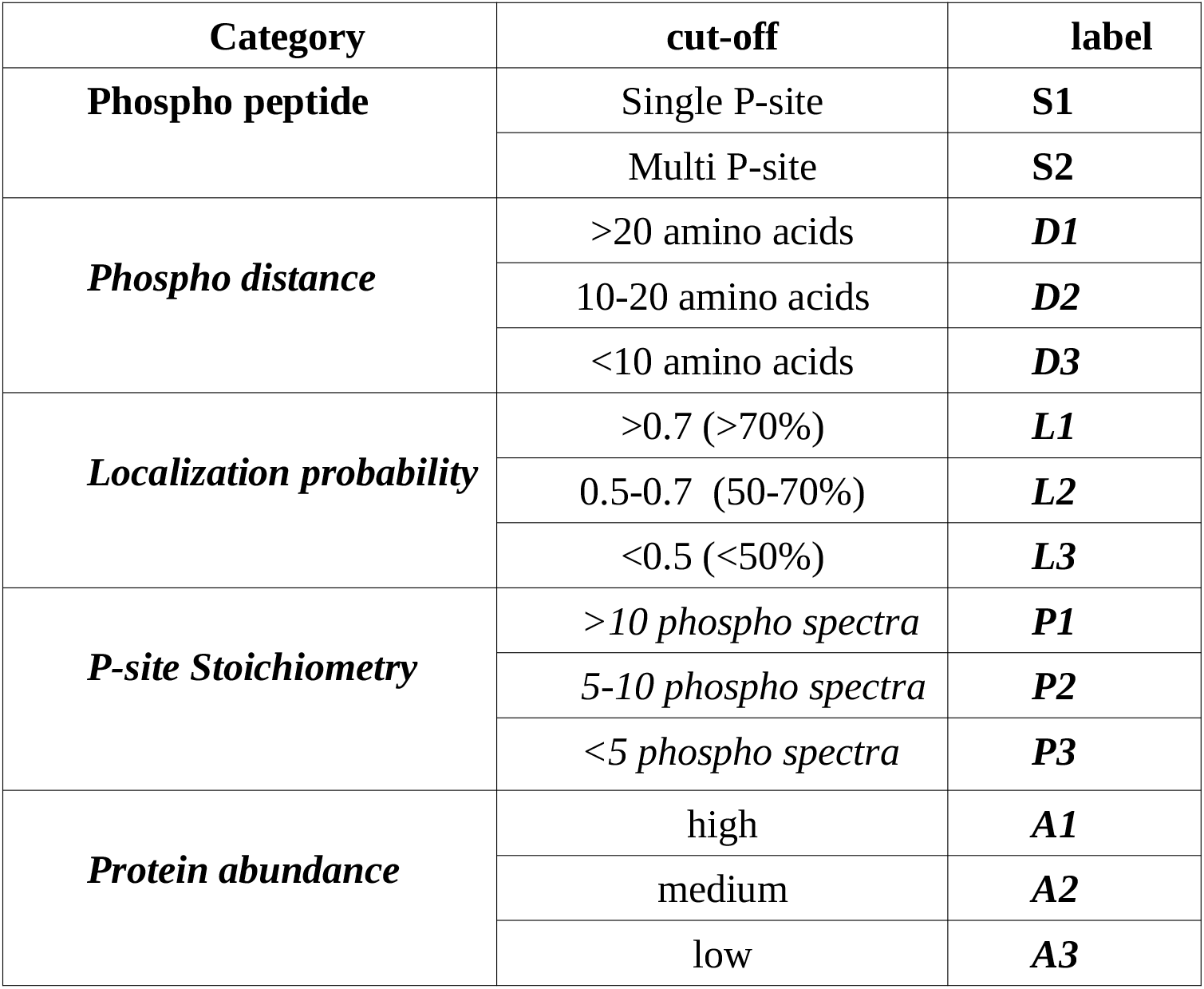
The five different meta-data at P-site level and the cut-off that were used to filter the reliable P-sites from the random ones.

The combined set of confident P-sites amounts to 81,404 (41,183 of which are MPS, and 40,221 are SPS), mapping to 12,632 phospho proteins (**Figure dataset overview, Supplementary Table T1)**. This set of phospho proteins and P-sites are considered the **positive set**.

To create a **negative set**, we retrieved all STY sites that do not have any evidence for phosphorylation in any of the four databases (Scop3P, UniProtKB/Swiss-Prot, PSP, and dbPTM) from the 12,632 proteins in the positive group. Random subsets were then drawn from these sites as needed to match the population size of the positive set for different downstream analyses (sequence, biophysics, and functional domains and structures).

### Structural proteins are highly phosphorylated in human samples

Almost 60% of all human proteins (12,632 proteins out of 20,365) were seen as phosphorylated in our dataset. 79% of these proteins were found to have less than 10% of their STY sites phosphorylated, while 21% had at least 10% of their STY sites phosphorylated (**Figure 3A, B and Figure 4A, B**). There are only very few proteins (41 proteins) which can be considered very highly phosphorylated (>=50%). Some of these proteins are structural proteins like prelamin A/C (P02545), keratin (P05787), histone H4 (P62805) and vimentin (P08670). Other proteins in this list include functional proteins like Leukemia-associated phosphoprotein (P16949), translation-initiation factor binding protein (Q13541), chromosome alignment-maintaining phosphoprotein (Q96JM3) and tumor protein (O43399).

**Figure 3:**
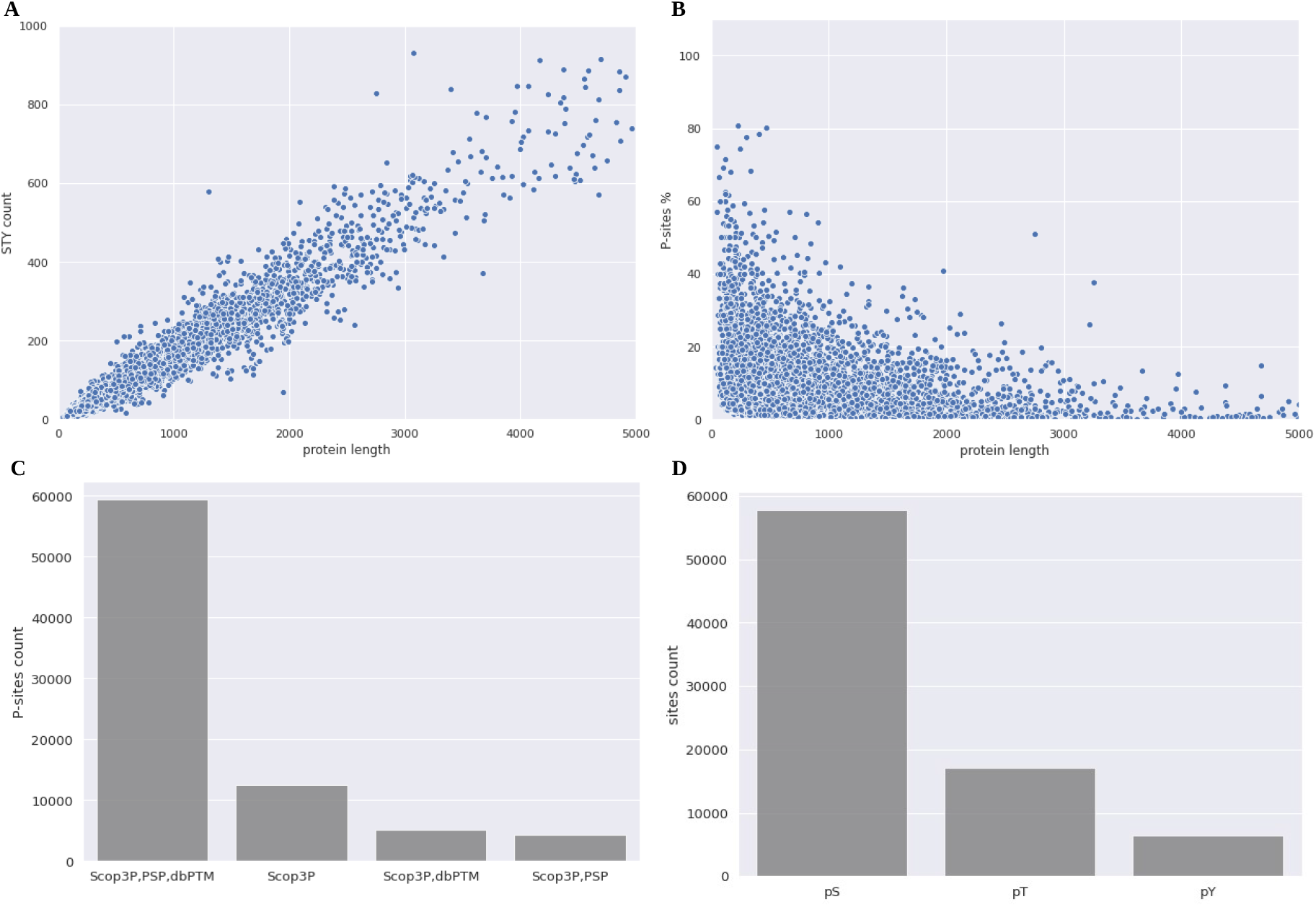
Final confident Phospho sites. **A)** Illustrating the relation between the total STY count to the phospho protein length (protein length in x-axis and STY count in y-axis: absolute counts are shown. Protein length restricted to 5000 amino acids in x-axis for better illustration). **B)** The relation between phospho protein length to the percentage of phosphorylation observed (protein length in x-axis and phosphorylation percentage in y-axis, protein length restricted to 5000 amino acids for better illustration) **C)** The distribution of final filtered P-sites (81,404) and the overlap between different re-sources and Scop3P. X-axis shows the absolute counts of P-sites in each sets and **D)** the distribution of total pS, pT and pY residues in final data (absolute counts are shown)

**Figure 4:**
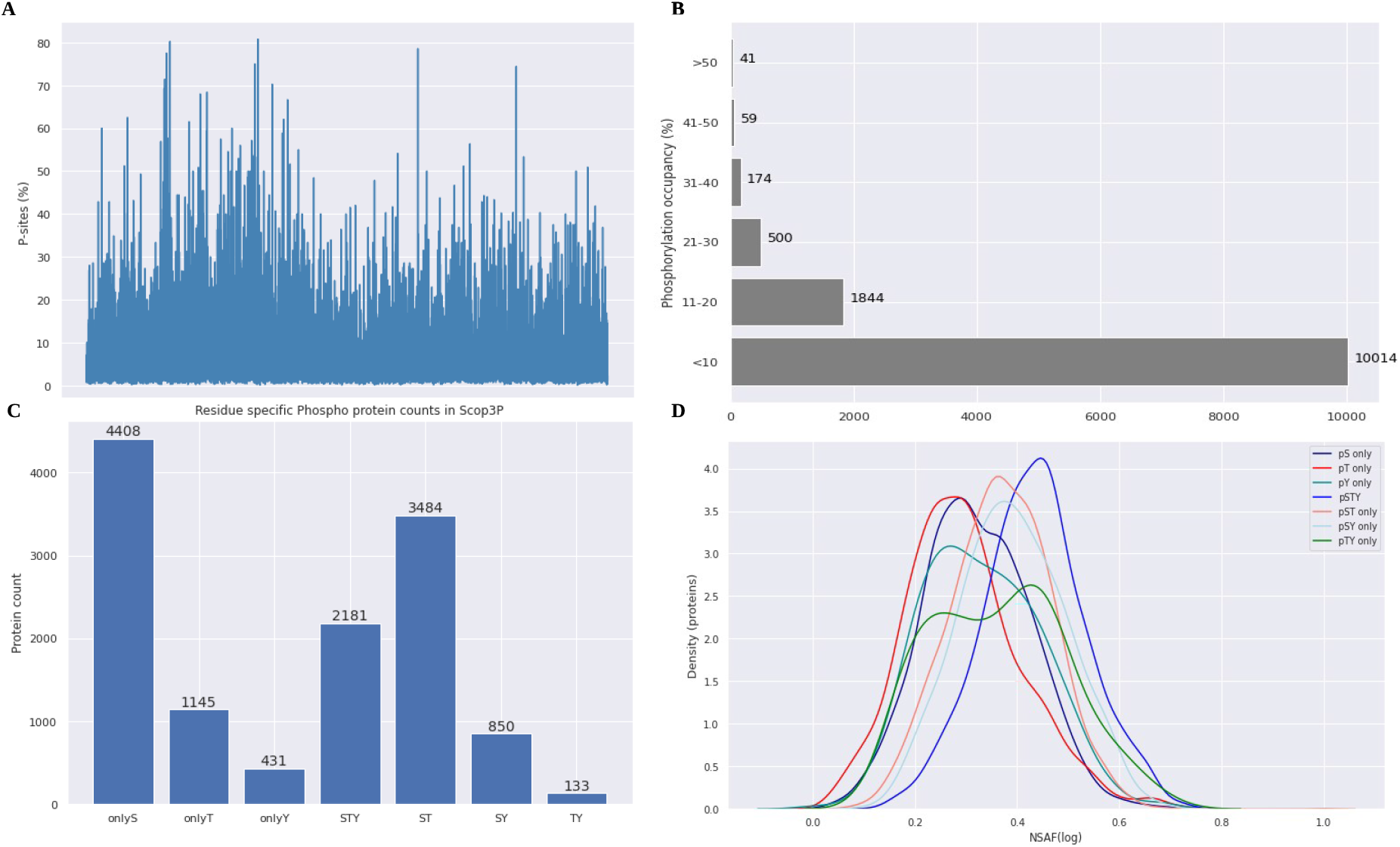
Phospho protein properties. **A)** The percentage of phosphorylation observed in proteins. x-axis denotes individual proteins **B)** The breakdown of number of proteins (x-axis) with at-least given percentage of phosphorylation (y-axis) **C)** Residue specific phospho proteins. Number of proteins seen as phosphorylated only in specific residues (S,T,Y) or in combination of different residues (ST, SY, TY) and proteins with all STY sites getting phosphorylated (x-axis is absolute count) and their NSAF abundance values **(D)** Proteins that are phosphorylated in different residues increases with increasing protein abundance.

As expected, the STY count increases with increasing protein length, but we observed an opposite trend in the phosphorylation percentage (**Figure 3A, B**). As a result, smaller proteins tend to show a higher phosphorylation percentage than longer ones (**Figure 3B**). 5.7% (pS:4%, pT:1.2%, pY: 0.5%) of all STY sites in the human proteome are found to be phosphorylated in our dataset. The distribution of the individual phospho residue types in our dataset is 71:21:8 (%) for pS, pT and pY respectively (**Figure 3D**). Although this aligns with previous findings that phosphorylation on Tyr (pY) occurs less often than pS/pT, if the relative amount of P-sites to the number of occurrences of their respective residue is considered, phosphorylation occurs on 3% of all tyrosines (Tyr) in the proteome, which is almost as frequent as Thr phosphorylation (3.6%), while Ser phosphorylation (7.8%) occurs twice as frequently.

Looking at residue specific phosphorylation coverage, we observed that there are proteins that have a tendency to be phosphorylated only on specific residues. For example, P60709 (beta-actin) has more observed pY; in this protein 33 (pS:12, pT:10, pY:11) out of 66 STY sites (S:25, T:26, Y:15) are phosphorylated. The overall phosphorylation percentage of this protein is 50%, but the individual residue percentage is higher for Tyr (~70%). Protein P62633 (cellular nucleic acid binding protein) is only phosphorylated on Ser and Tyr residues (pS:6, pY:2). We therefore classified the phospho proteins in our dataset in seven different groups depending on the residue type(s) which are found to be phosphorylated (onlyS, onlyT, onlyY, STY, ST, SY, and TY) (**Figure 4C**). The largest group is that of proteins only phosphorylated on Ser (S), followed by proteins phosphorylated on both Ser and Thr (ST) residues, with proteins phosphorylated on all three residues (STY) in third place. Interestingly the protein group phosphorylated on “only Y” contains 431 proteins that have only Tyr phosphorylation observed, despite these proteins containing more Ser and Thr residues than Tyr. This might indicate very specific phosphorylation targeting on these proteins.

We checked whether there is any relation to protein abundance and a residue specific nature of phosphorylation, which revealed an interesting trend: the number of different sites that are phosphorylated in a protein increases with protein abundance. For example, proteins with all three sites (STY) phosphorylated tend to be highly abundant, while phosphorylation of pST and pTY tends to occur in moderately abundant proteins. Interestingly, proteins containing only pT residues tend to be of lower abundance, followed by onlyS, onlyY and onlyTY phospho proteins. (**Figure 4D**).

### Distribution of P-sites over normalized protein sequence length

To address the question whether phosphorylation might be more frequent in certain regions of a protein, such as their N- or C-terminal regions, we mapped all proteins to a normalized sequence length of 1-100 and re-scaled their phosphorylation positions within this range (**Figure 5**). The distribution of P-sites on this normalized sequence length scale shows that P-sites are seen in almost all regions of human proteins (**Figure 5**) with slightly more P-sites seen in the C-terminal region.

**Figure 5:**
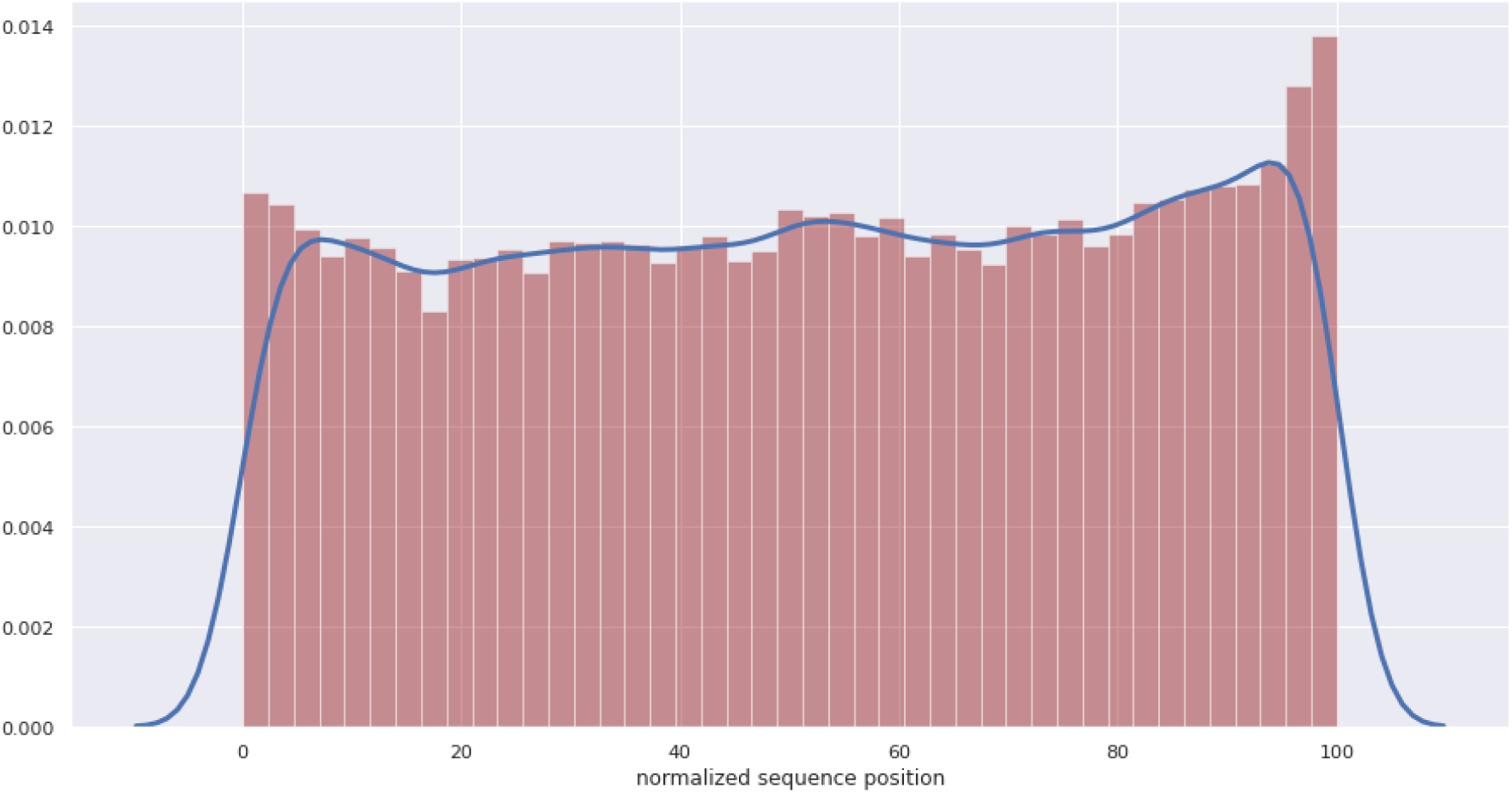
P-sites distribution on a normalized protein sequence. A normalized protein sequence mapped with P-sites showing where the protein is most phosphorylated in the proteome. All proteins of different lengths were scaled to a uniform length of 100 amino acids and the identified P-sites locations were re-scaled to fit within this range. In contrary to previous findings that P-sites are predominant in terminal regions, we see P-sites are distributed across the protein length. There is a slight increase in the C-terminal region of proteins than the N terminal. x-axis shows the scaled position 1-100 amino acids and y-axis the distribution of P-sites

### Phosphorylation in functional domains

To investigate the potential functional impact of protein phosphorylation, we explored the protein domain-context of phosphorylation. P-sites were matched to the Pfam^53^ domain boundaries for the respective protein. We observed that 25% (of which, pS: 16%, pT: 6%, pY: 3%) of all P-sites (20,954 out of 81,404) occur in Pfam domain regions, constituting 3,537 different functional domains in 7,003 proteins.

We looked at the number of different protein instances that carry a specific domain to find the most frequently represented domain families in our dataset (**Figure 6A**) shows the top domains, which are found in 25 or more proteins; **Supplementary Table T2** provides the complete data). As expected, we found domains involved in signaling and regulation (kinase, RAS, PH, PDZ) to be most frequently represented, followed by structural domains (actin, myosin, histone). Interestingly, we also find a lot of RNA recognition motif (RRM) domains to be phosphorylated, and this across 98 different protein instances (**Figure 6A**).

**Figure 6:**
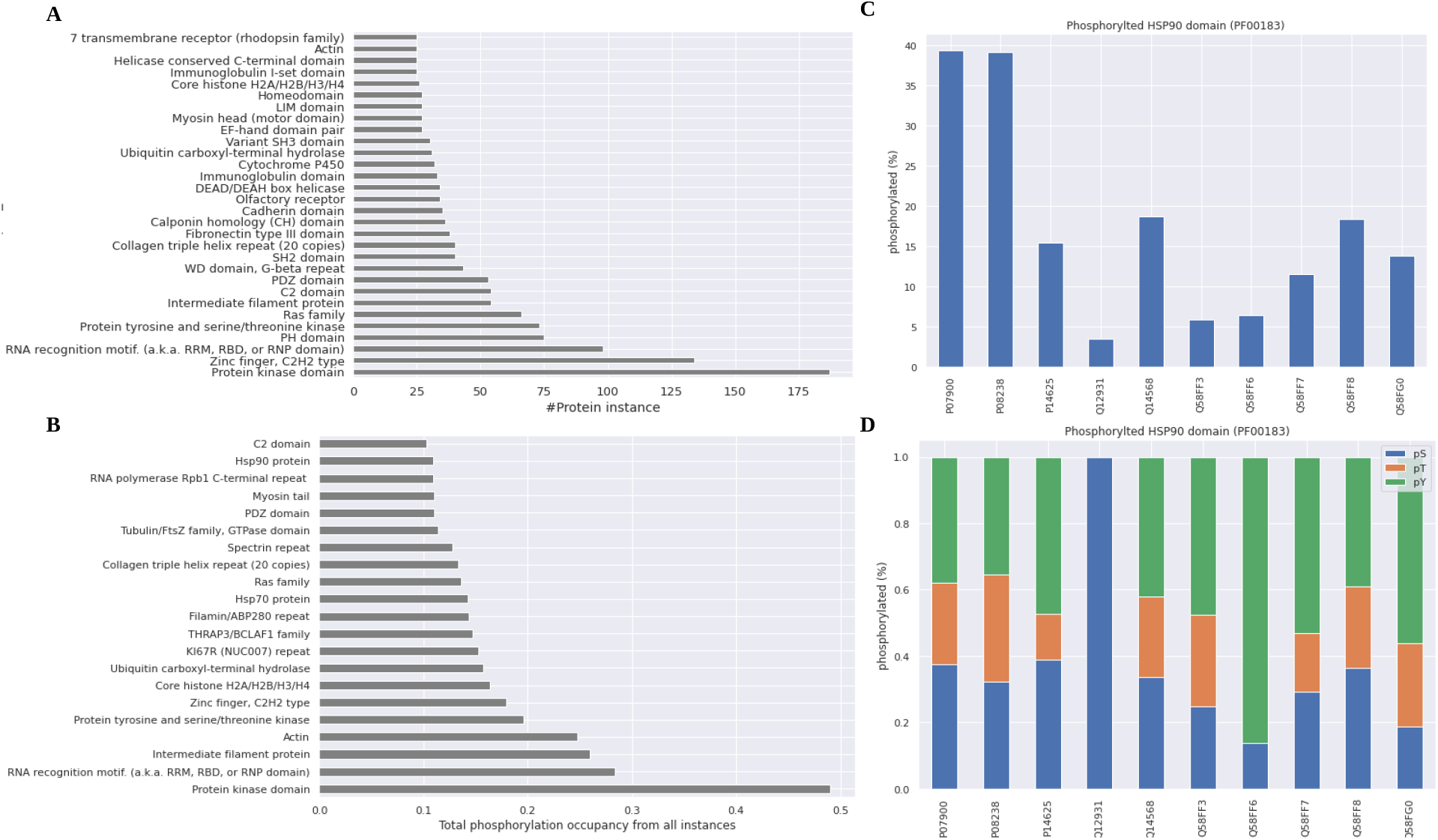
Phosphorylation on functional domains. **A)** Number of different protein instances (x-axis absolute count) a domain is seen as phosphorylated (data shown only for domains with instances >=25). Kinase being one of the most abundant protein in human proteome, kinase domain is seen as phosphorylated in 175 instance followed by domains involved in interactions like Zinc finger, RRM, PDZ and PH domains **B)** The average phosphorylation observed by pooling all instance of the domain together. Actin which has less number of instance (25 proteins as shown in A) but the overall phosphorylation status is higher than zinc finger C2H2 domain with more than 125 instance. The percentage of actin getting phosphorylated is higher than zinc finger, C2H2 irrespective of the number of instance. x-axis shows the average percentage of phosphorylation obtained after pooling all instance. **C,D)** The difference in phosphorylation seen in different protein instances (x-axis) of the same domain (HSP90) because of difference in sequence composition in protein **(C)** and the type of amino acid phosphorylated in different instance **(D)** Different protein instance of the domain is shown is x-axis ((UniProt protein ID) and the percentage of phosphorylation on y-axis

We then estimated the average phosphorylation occupancy of a domain (**Supplementary Table T3**) and found that 35% of all domains can have at least 10% of their STYs phosphorylated. Looking at the overall contribution of the phosphorylations found in a given domain family to the total number of P-sites in our dataset (81,404 P-sites) (**Figure 6A**), we calculate the relative P-site contribution of each domain as the number of P-sites in that domain divided by the total number of P-sites in our dataset. This shows that nearly half of all P-sites are derived from kinase domains, but that domain families with fewer protein instances (histone core, HSP70 and HSP90) contribute more phosphorylations than domain families with more protein instances (PDZ, RAS, PH, C2) (**Figure 6B**).

Because domain boundaries and sequence composition vary between proteins, the phosphorylation occupancy may differ for the same domain between different protein instance, and which residues (S, T, or Y) are observed to be preferentially phosphorylated. We therefore analyzed occupancy and residue preference for two domains with several protein instances: HSP90 (PF00183) and HSP70 (PF00012).

For the HSP90 (PF00183) domain, which is involved in various functions including protein folding, protein heat stability, and protein degradation, is seen as phosphorylated in ten different proteins in our dataset, but the phosphorylation occupancy can differ substantially between these different proteins (**Figure 6C**). HSP90 alpha (P07900) and HSP90 beta (P08238) are both localized within the nucleus and show very similar occupancy (**Figure 6C**) as well as very similar residue preference (**Figure 6D**). On the other hand, endoplasmic HSP90 (P14625) and mitochondrial HSP90 (Q12931) both have low occupancies, and quite distinct residue preferences. Endoplasmic HSP90 has more pY sites and less pT sites when compared to nuclear HSP90 alpha and beta, while mitochondrial HSP90 has only pS sites (**Figure 6D**).

Similar observations can be made for HSP70 (PF00012), which is also involved in protein folding, and is seen as phosphorylated in fifteen different proteins in our dataset (**Supplementary Figure S3**). Proteins O95757, P38646 and P11142 are annotated as localized in nucleus/nucleolus and have similar occupancy, and all involve pS, pT and pY phosphorylation (**Supplementary Figure S3**). Protein P54652 is localized in spindle, and has no observed pS phosphorylation, while Protein P48723 is localized in the microsome/endoplasmic reticulum (ER) and has no observed pY sites (**Supplementary Figure S3**). Interestingly, protein P11021 is also annotated as localized in ER but specifically in the ER lumen and has evidence of pS, pT and pY in HSP70 domains.

Based on these observations of differing occupancies and observed residue preferences for a given domain across different proteins, we can conclude that we need to take this diversity into account when attempting to generalize for a given domain.

We can classify all domains in our dataset based on the residue type(s) that are being phosphorylated across all their protein instances: onlyS, onlyT, onlyY, STY, ST, SY and TY (**Supplementary Figure S4:A**). For example, Histone deacetylase (PF00850) is seen in four instances (O15379, Q8WUI4, Q96DB2, Q9UKV0) and is always seen phosphorylated on pS only (**Supplementary Figure S4:B**), resulting in an onlyS classification. On the other hand, zinc binding dehydrogenase (PF00107) is seen in nine instances with either pT, pS or pST phosphorylation, so this domain is classified as “ST” (**Supplementary Figure S4:C**), and actin domain (PF00022) has 25 instances, with phosphorylated STYs in several instances, leading to “STY” classification (**Supplementary Figure S4:D**). We find that most domains (30%) are seen as onlyS, followed by ST, STY, SY, onlyT, onlyY, and onlyTY phospho domains (**Supplementary Figure S4:A**).

We also found that there are 2,761 proteins in our dataset where phosphorylation occurs only on their functional domains. These include single domain proteins where the domain spans the length of the protein, such as beta-actin (P60709) and transcription factor GTF2F1 (P35269), as well as multi domain proteins (Q92997, P15311, P54753) where evidence of phosphorylation is seen in more than one of their domains.

### Disorder properties of P-sites

We could annotate 11,880 of the 12,632 proteins in our dataset with disordered region information. Most phosphoproteins have a low fraction of their sequence annotated as disordered, with only 21% of phosphoproteins having more than 50% of their sequence annotated as disordered (**Figure 7A**). 132 proteins are reported to be completely (100%) disordered (**Figure 7A**). After mapping P-sites to disordered regions, we found that 65% (53,276 sites) of all P-sites (from 8,171 proteins) are located in intrinsically disordered regions or in regions involved in structural transitions and/or phase separation (**Figure 7D)**. Moreover, 3,709 phosphoproteins do contain annotated disordered regions, but none of the identified P-sites in these proteins (7,991 P-sites) are present in their disordered regions. It should be noted that the disordered fraction for these 3,709 proteins also tends to be very low i.e., they appear to be primarily ordered (**Figure 7B**). We could not find any disorder annotation for 752 proteins (containing 2,486 P-sites).

**Figure 7:**
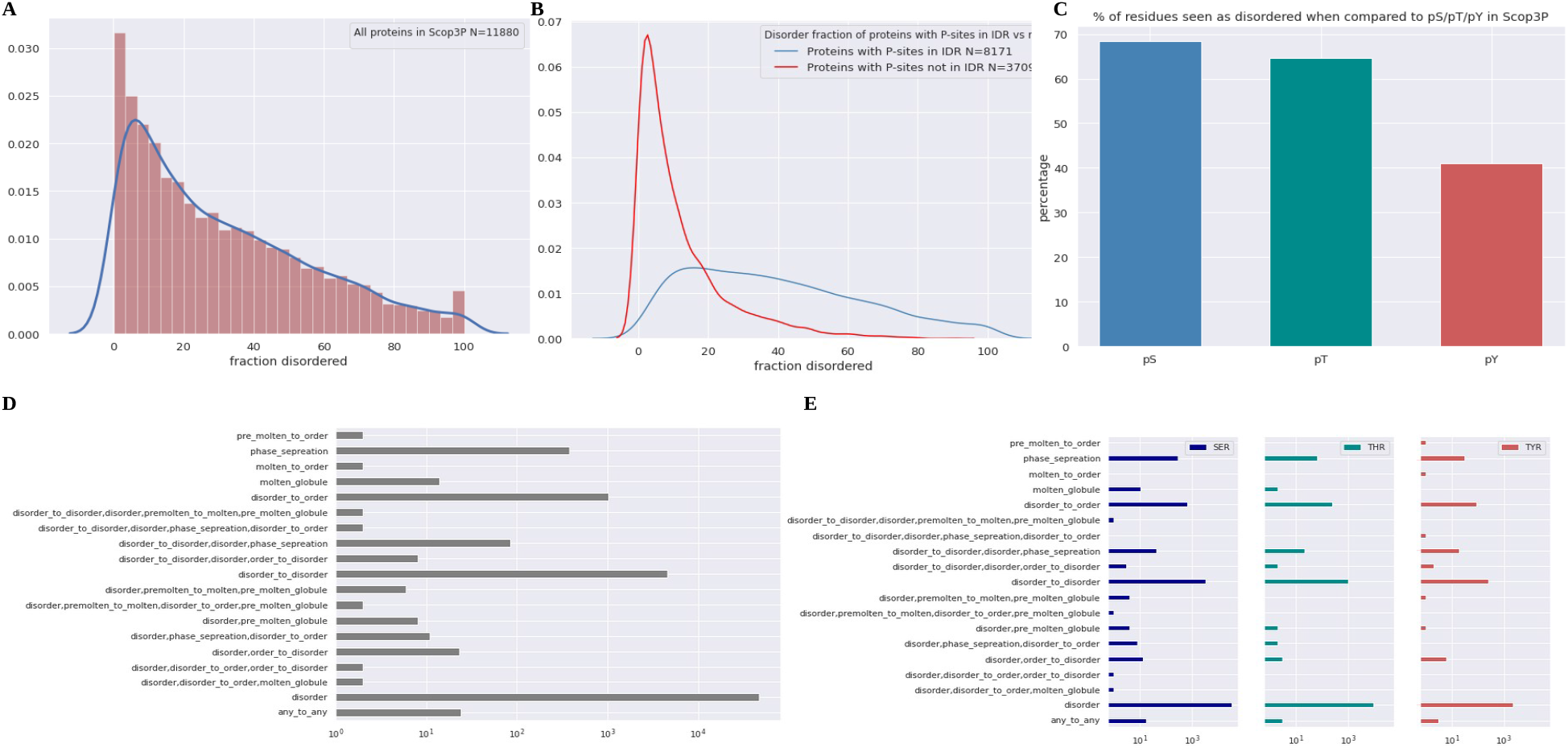
Disorder and structural transition properties of phospho proteins and P-sites. **A)** Shows the fraction of disorderness (x-axis) in the protein and the number of proteins in that fraction (y-axis) **B**) Disordered fraction (x-axis) of proteins that contain P-sites in disordered regions (blue) and the proteins with all P-sites that are not in disordered regions (red). y-axis shows the density of proteins in that fraction **C)** Distribution of percentage of pS, pT and pY residues in disordered regions **D)** Distribution of P-sites seen in different structural transition states. The structural states are shown in y-axis and the P-sites count (log transformed) in x-axis **E)** Distribution of number of residues pS/pT./pY (x-axis (log transformed) that are observed in different structural transition states (y-axis).

The fraction of pS, pT and pY in disordered regions over the total number of pS, pT, and pY sites shows that almost 70% of all pS, over 60% of all pT sites, and 40% of all pY sites are seen in disordered regions/states (**Figure 7C**). Interestingly, most of the P-sites are in disorder regions, or in transition regions involving disorder (**Figure 7D**). As to individual sites, pS sites are seen most frequently, and occur in almost all transition states, while pT sites are not seen in some states like ‘molten globule’ and ‘pre-molten to molten’ transitions. pY sites on the other hand, are seen in ‘pre-molten to order’ and ‘molten to order’ states that are not seen in pT (**Figure 7E**). Nevertheless, the overall distribution of pT and pY sites across states resembles the one for pS sites.

### Structural properties of P-sites

It has been shown previously that phosphorylation often occurs in the disordered/flexible regions of proteins^20,21^. This is because these sites must be accessible to kinases, phosphatases or regulatory enzymes that recognize these sites. But some P-sites are found buried in solvent inaccessible regions of proteins as well^21,68^. To better understand how P-sites in solvent inaccessible regions are phosphorylated, we analyzed the sites in different structural context with specific focus on possible allosteric mechanisms like a conformational switch that exposes such sites and makes it accessible for kinase.

After mapping P-sites to the experimental 3D structural coordinates and filtering (**See methods**) we find that 10% of all P-sites in our dataset can be annotated with structural information (8,432 P-sites from 3,059 proteins). Of these structurally annotated P-sites, 58% were pS, 25% pT, and 17% pY sites. We also found a few P-sites with the phospho-group attached in the structure (SEP (pS), TPO (pT) and PTR (pY), with 25, 28, and 23 occurrences, respectively, corresponding to ~0.3% of all structurally annotated P-sites each). While most P-sites with structural annotation (54%) are seen in flexible regions with coil propensity, we also find a considerable number of P-sites (40%) in structured regions with helical or strand propensity. The remaining 6% of P-sites were found to be in regions missing from the structure (as these regions are typically disordered and not visible by X-ray crystallography), or without any structural propensity annotation in DSSP (**Figure 8A ii**). The distribution of secondary structural propensity for individual residue types **(Table 2)** shows that most of the pS and pT residues are in unstructured/coil regions, with pT residues favouring beta strand propensity more than pS residues, while pY sites are seen more frequently in structured regions of proteins (55.2% sites combined in helix and strand regions) (**Table 2**).

**Figure 8:**
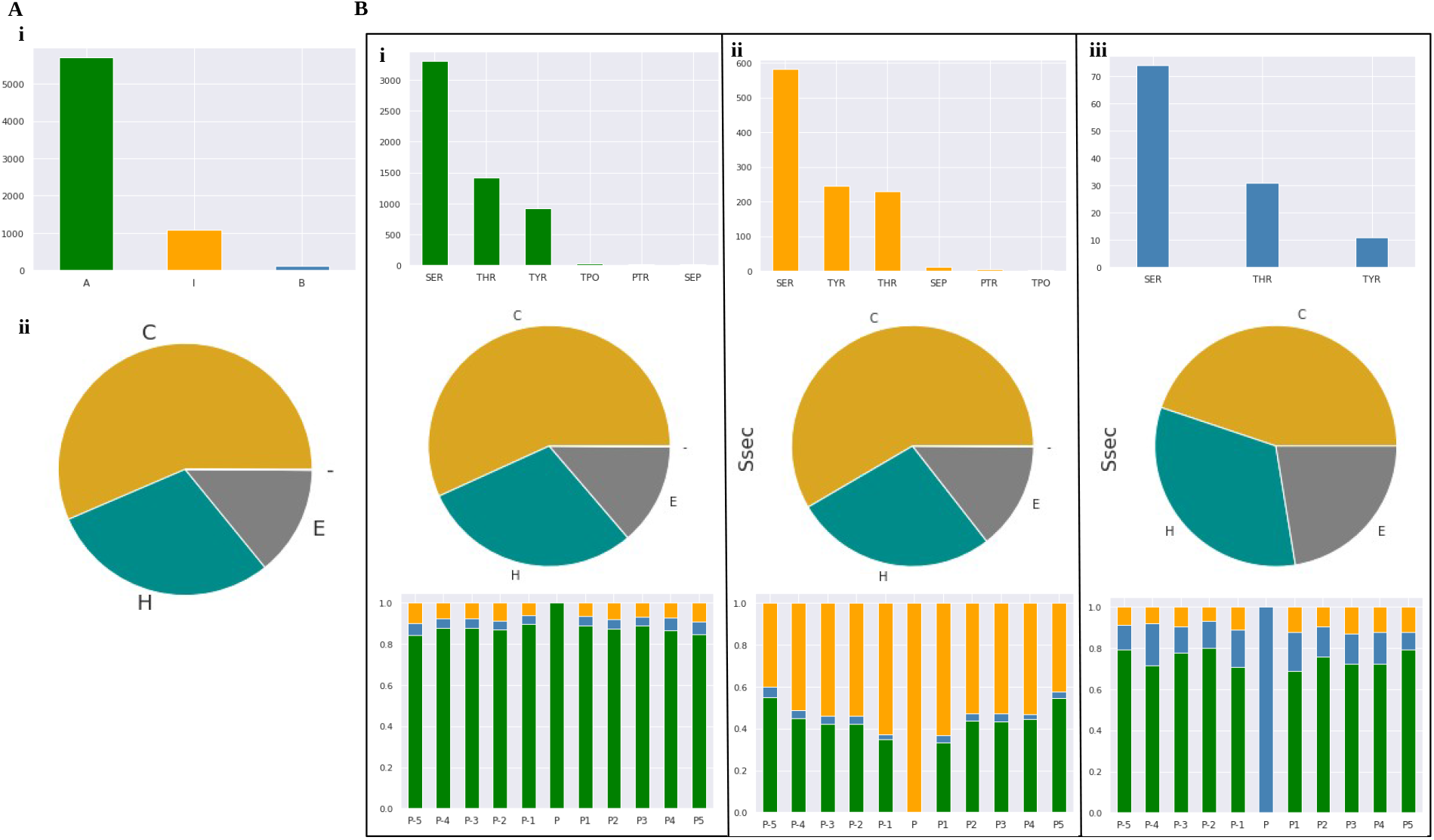
Structural properties of P-sites. **A) i)** Distribution of P-sites seen in solvent accessible (**A**-green), interface (**I**-orange) and in buried regions (**B**-blue). And **(ii)** their secondary structural states (**H**: Helix, **C**: coils/unstructured, **E**: beta strand, -: not available/missing regions) y-axis shows the absolute count. **B)** The distribution of amino acid types in accessible, interface or buried regions (**i, ii, iii: Top panel**, absolute counts in y-axis) and the breakdown of secondary structural properties in accessible, interface and buried regions is shown (**i, ii, iii: middle panel**). The bottom panel shows the solvent accessibility of flanking 5 residues in N (P-5 to P-1) and C terminal (P1 to P5) around the central P-sites (P) in accessible, interface and buried regions. y-axis shows the log transformed absolute counts.

**Table 2:** Distribution of secondary structural propensity of individual residue types

|  | pS (%) | pT (%) | pY (%) |
| --- | --- | --- | --- |
| coil/unstructured (C) | 60.0 | 56.3 | 36.5 |
| helix (H) | 24.0 | 23.5 | 33.0 |
| beta strands (E) | 9.0 | 14.7 | 22.2 |
| missing segments (--) | 6.0 | 5.3 | 8.3 |

Because secondary structures only tell us the propensity of the residue to form a structural element, but does not address how accessible that residue’s side chain is, we looked at the solvent accessibility of these residues specifically. We performed this accessibility analysis on 6,893 P-sites for which this information was available, out of 8,432 total P-sites. Most of these sites (5,701) are indeed found in solvent accessible regions, while 1,076 are in interface regions, and only 116 are found in buried regions (**Figure 8A i**). The residues which are also seen as phosphorylated in structures (SEP, PTR and TPO) were never observed in buried regions (**Figure 8 B i, ii, iii top and middle panel**).

We also analyzed the accessibility of flanking residues, i.e., five amino acids before and after each P-site in the PDB sequence. Flanking residues around solvent accessible P-sites are also seen as accessible, while flanking residues around interface P-sites tend to be in interface regions as well. Interestingly, flanking residues around buried P-sites tend to be far more accessible than the P-site itself (**Figure 8B i, ii, iii bottom panel**).

### Mutation hot-spots on and around P-sites

We investigated potential mutational hot-spots by analyzing the frequency of mutations that occur on and around P-sites. We mapped the sequence positions of 79,205 natural variants obtained from Humsavar to the 81,404 filtered P-sites in our dataset. A total of 5,034 mapped mutations occurs either directly on P-sites or in flanking sites (± 5 residues N- and C-terminal to the P-site). We classified these mutations into three different classes: (i) ‘direct’ (mutations that occur on P-sites), (ii) ‘proximal’ (within two residues to P-sites), and (iii) ‘distant’ (between three and five residues from the P-site). After filtering for flanking five amino acids, our final data contains 4,653 mapped mutations: 395 mutations are ‘direct’, 1,692 are ‘proximal’, and 2,566 are ‘distant’ mutants. Mutations that occur directly on P-sites tend to be neutral rather than harmful (**Figure 9A**); only 134 (pS:68, pT:36, pY:30) out of 395 direct mutants were labeled as disease causing.

**Figure 9:**
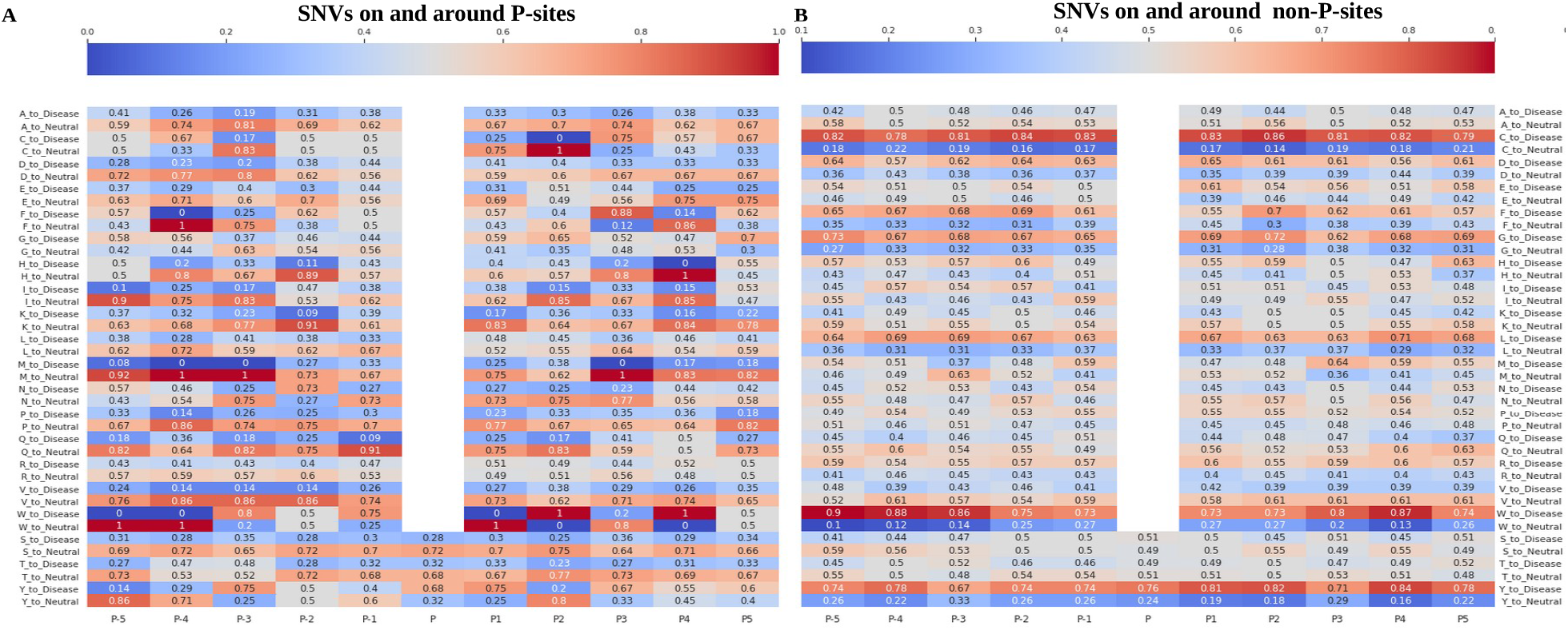
Mutational hot-spots on and around P-sites and non P-sites. Illustrates the frequency of amino acid variants in central (P) and its flanking N (P-5 to P-1) and C terminal (P1-P5) sites around P-site (**A: left panel**) and non P-sites (**B: right panel**). The absolute counts are normalized on columns and the normalized values are shown. The color range from blue (less frequent) to red (more frequent). x axis shows the location of mutation. P denotes the mutation is seen is central site (phospho STY/non-phospho STY) and P1 means there is a mutation next to a P-site in the C terminal position and P-5 denotes a mutation at 5^th^ residue in the N-termini when there is P-site at position ‘P’. Y axis shows the variant amino acid type for every position followed by if the variant is disease causing or neutral. Every cell in the plot represents the frequency of mutation seen at that position.

We looked at the frequency of amino acids that are mutated in different phospho residues (pS, pT and pY). The disease-causing mutants on pS are more varied (Y, W, P, C, A, T, N, R, I, F, L) than pT (K, A, P, N, R, I, S, M) and pY variants (C, H, N, D, S, F). The most frequent variants for pS (F, P, L) and pT (I, A, M) tend to be more hydrophobic while the most frequent pY variants are more polar and basic amino acids (C, S, H).

We then looked at the mutational hot-spots around the non-P-sites to see how often a non-P-site and/or the residue flanking the non-P-sites are being mutated when compared to its phosphorylated counterparts, including their frequency of being disease causing or neutral polymorphism (**Figure 9A and B**). The direct mutations on P-sites tend to be more neutral polymorphic, except for pY variants (**Figure 9A**), which tend to be more disease causing. For example, the frequency of Tyr (Y) mutation at position P2 causing deleterious effects are much lower (**Y_to_Disease: 0.2)** when there is a P-site at position P and in contrary for the same position P2, a Tyr mutation is more deleterious if the site P is not phosphorylated (**Y_to_Disease: 0.82**).

The proximal and distant mutations, when there is a pS or pT at central site, tend to be more neutral. Interestingly, for pY variants, the proximal mutants at P1 tend to be more disease causing while P-1 is more neutral. There is a slight increase in frequency of disease-causing variants at flanking regions P-3, P3-P5 (**Figure 9A**). For the non-phosphorylated central sites, non-phospho Ser (S) and non-phospho Thr (T) shows ambiguous patterns on and around flanking sites, while non-phospho Tyr (Y) tends to be more disease causing irrespective of the location (**Figure 9B**).

We saw some interesting signals among the mutations observed in flanking sites. The mutation on Trp (W) residue is disease causing at any location around the non-P-sites and are disease causing at P-1, and P-3 around P-sites (**Figure 9A,B**). The variation on Arg (R) residue is ambiguous around the P-sites, the N-terminal sites were more disease causing and in C-terminal there was not much difference, while around non-P-sites, it appears to be more deleterious. Pro (P) variation tends to be neutral around P-sites and ambiguous around the non-P-sites. The frequency of Pro variation to be neutral is high around P-site than non-P-sites. This is interesting because many kinases prefer a Pro at P1 position for effective binding^15,69^ and the mutation at Pro does not make any difference on the phenotype in our analysis. Mutations on Phe (F) were more disease causing around non-P-sites and around P-sites they were more neutral except P-2, P3 and P5 which are more deleterious. Variations on Leu (L) are more disease causing around non-P-sites while neutral around P-sites. Gly (G) variants are more deleterious around non-P-sites and are more neutral around P-sites except the proximal P2 and distal P5 sites, which are disease causing (**Figure 9A, B**). Cys (C) variants are more disease causing at P-4, P3, P4 and P5 sites. Variations on Ala (A) residue are more neutral around P-sites and are ambiguous around non-P-sites. Asp (D) residues were more neutral around P-sites and disease causing around non-P-sites. Variations on Val (V) residues are neutral irrespective of the location and phosphorylation status. Other variants like Ile (I), Lys (K), Gln (Q), Glu (E) are more neutral around P-sites and ambiguous around non-P-sites except for Glu (E) which is more disease causing at P1 (**Figure 9A, B**).

The above analysis shows the Humsavar-annotated variation of wild type amino acids on and around P-sites to any possible destination residue. We also looked for STY directed mutations i.e., how often a P-site or a residue in its flanking region is mutated to Ser/Thr/Tyr, as these create possible new P-sites (**Supplementary Figure S5**). Our data shows that a Ser residue is mutated to Tyr or Thr, while Tyr and Thr is only mutated to Ser, irrespective of their phosphorylation status. The frequency of ser mutation is more at the central site (P) and is more disease causing when its mutated to Tyr in the non-phospho context. Tyr that is mutated to Ser in both phospho and non-phospho context tends to be more disease causing while Thr mutation to Ser is more nutral.

### Biophysical properties of phospho variants

We looked at the biophysical properties of wild type primary protein sequence to understand if there is any difference in the dynamics that favors mutation at pSTY sites. Moreover, we also investigate whether there is any difference between neutral and harmful variants. In total, our data contains 389 pSAVs (mutations occurring at P-sites) and 79,606 variants that are not in P-sites. First we looked at the biophysical properties of pSAVs that are disease causing to the ones that are neutral. Sites that are disease causing tend to be more rigid than neutral polymorphic sites (**Figure 10 A, B**). Disease causing P-sites have a more rigid backbone (p-value: 7.24 x 10^-10^) (**Figure 10A**), less disorder propensity (p-value: 6.65x 10^-6^), increased side chain dynamics (p-value: 5.87x 10^-6^), and also tend to have higher early folding propensity^60^ (p-value: 8.23x 10^-5^) (**Figure 10B, C, D**) as compared to neutral polymorphic sites.

**Figure 10:**
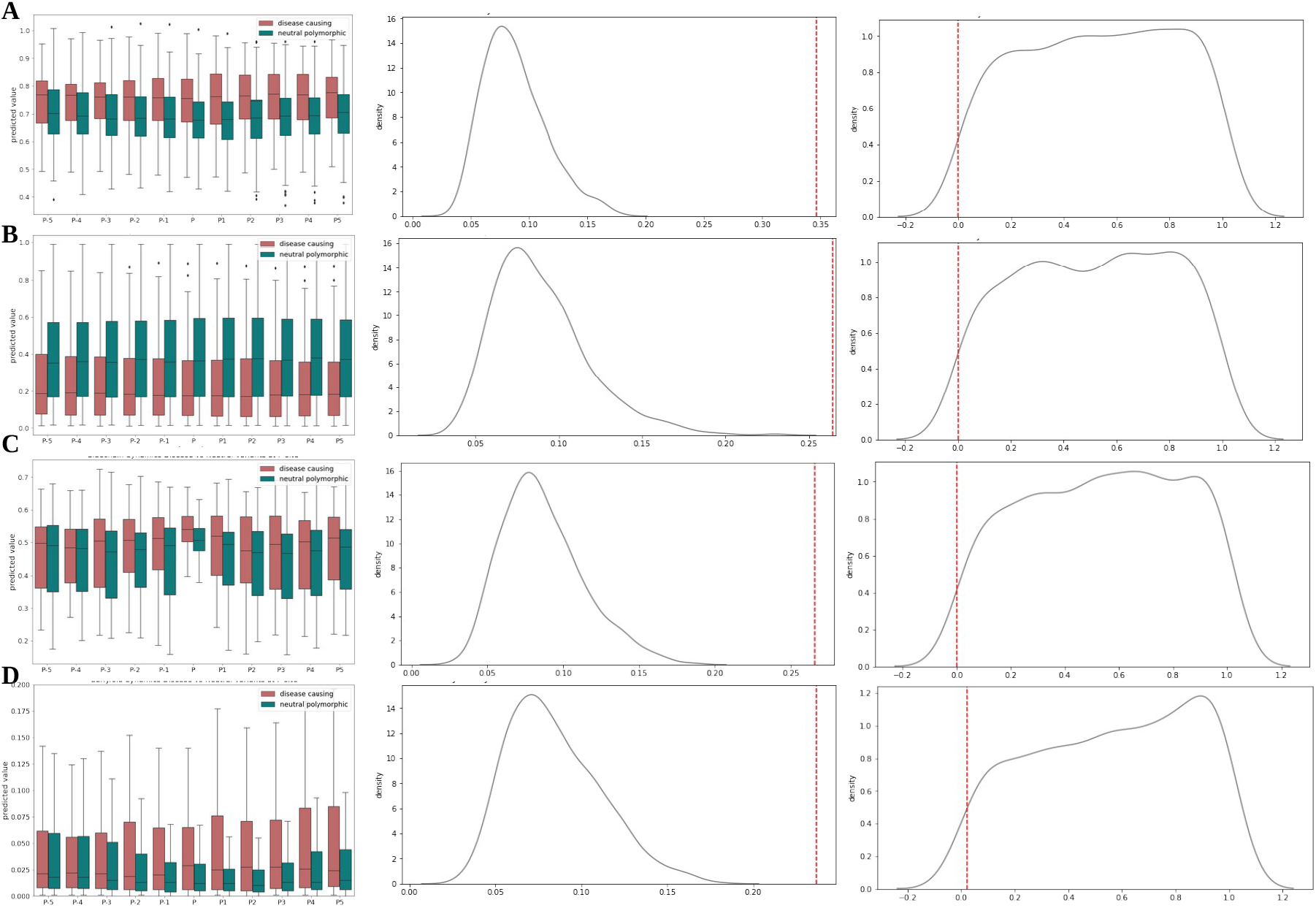
Biophysical properties of Disease causing and neutral phopsho variants. **Left panel** shows the distribution of dynamic values of disease causing phopsho (Indian red, N=134) and neutral phospho variants (dark cyan, N=255). **Middle panel** shows the KS-Dstat distribution of 1000 random permutation test by shuffling phopsho variants and neutral variants and sampling (size=389: red line is the actual observed value between two group) and **right panel** shows the P-value distribution of random permutation test (red line is the actual observed value). Predictions are done on primary amino acid sequence. Plot shows the biophysical preference for disease causing and neutral phospho variants for central site (P) and its flanking sites (P-5 to P5). KS statistics are shown only for the data points at central site (position ‘P’) **A)** Distribution of Backbone dynamic values. Values > 0.8: rigid, 0.69-0.8: context dependent, <0.69: flexible. **B)** Distribution of disorder propensity values. Values above 0.5indicate that this is likely a disordered residue. **C)** Sidechain dynamics: Note at ‘P’ its just S/T/Y residues and the flanking residues can be any amino acid. Higher values mean more likely rigid. These values are highly dependent on the amino acid type (i.e. a Trp will be rigid, an Asp flexible). **D)** Early folding propensity. Values above 0.169 indicate residues that are likely to start the protein folding process, based on only local interactions with other amino acids.

Then we compared the mutations at P-sites to the ones that are not in P-sites. Direct mutants tend to be more rigid (**Supplementary Figure S6:A, B**) with rigid backbone dynamics (p-value: 6.12 × 10^-5^) (**Supplementary Figure S6:A**) and increased side chain dynamics (p-value: 5.19 × 10^-5^) (**Supplementary Figure S6:C**). The disorder and early folding propensities also show a difference with less disordered and increased early folding propensity for direct mutants (p-value: 0.002 for disorder propensity and p-value: 0.02 for early folding) (**Supplementary Figure S6:B, D**).

### Biophysical properties of P-sites

Not all STY sites in human proteins are phosphorylated, even though most of these non-P-sites are in solvent accessible regions. To understand why only some of these solvent accessible sites are phosphorylated, and, conversely, how some solvent inaccessible/buried sites are phosphorylated, we looked at several biophysical properties: backbone and side chain dynamics, disorder propensity, and early folding propensity of both P-sites and non-P-sites. We performed the analysis on different sub-groups, based on their known functional or structural annotation (**Figure dataset overview**):

1. Properties of P-sites and non-P-sites at primary amino acid sequence level
2. Properties of P-sites and non-P-sites in functional domains
3. Properties of P-sites and non-P-sites with three-dimensional structural information

We calculated the KS test statistic against a null distribution estimated by random sampling without replacement from the negative set to match the population size of the positive data (**See Methods**).

### Biophysical properties at sequence level

The biophysical properties were predicted on the amino acid sequences of the proteins. Because we wanted to compare the flanking ± 5 residues we considered only sites that have 5 flanking residues on both sides. After this filtering, the final positive set (P-sites) contains 79,994 sites, and the negative set (non-P-sites) has 1,116,380 sites (from the same proteins that are in the positive set) (**Figure dataset overview**).

#### STY sites that are phosphorylated tend to be more dynamic and flexible

pSTY sites tend to be more flexible than non-phosphorylated sites (**Figure 11**). P-sites are more dynamic in their backbone and highly disordered when compared to non-P-sites (**Figure 11A, B**). P-sites not only have a flexible backbone, but their side chains are predicted to be more flexible as well (**Figure 11C**). P-sites have lower early folding propensities^60^ as compared to non-P-sites, indicating that most of the STY residues involved in so-called foldon (protein region that folds as a unit) formation tend not to be phosphorylated (**Figure 11D**), which fits with their tendency to be buried.

**Figure 11:**
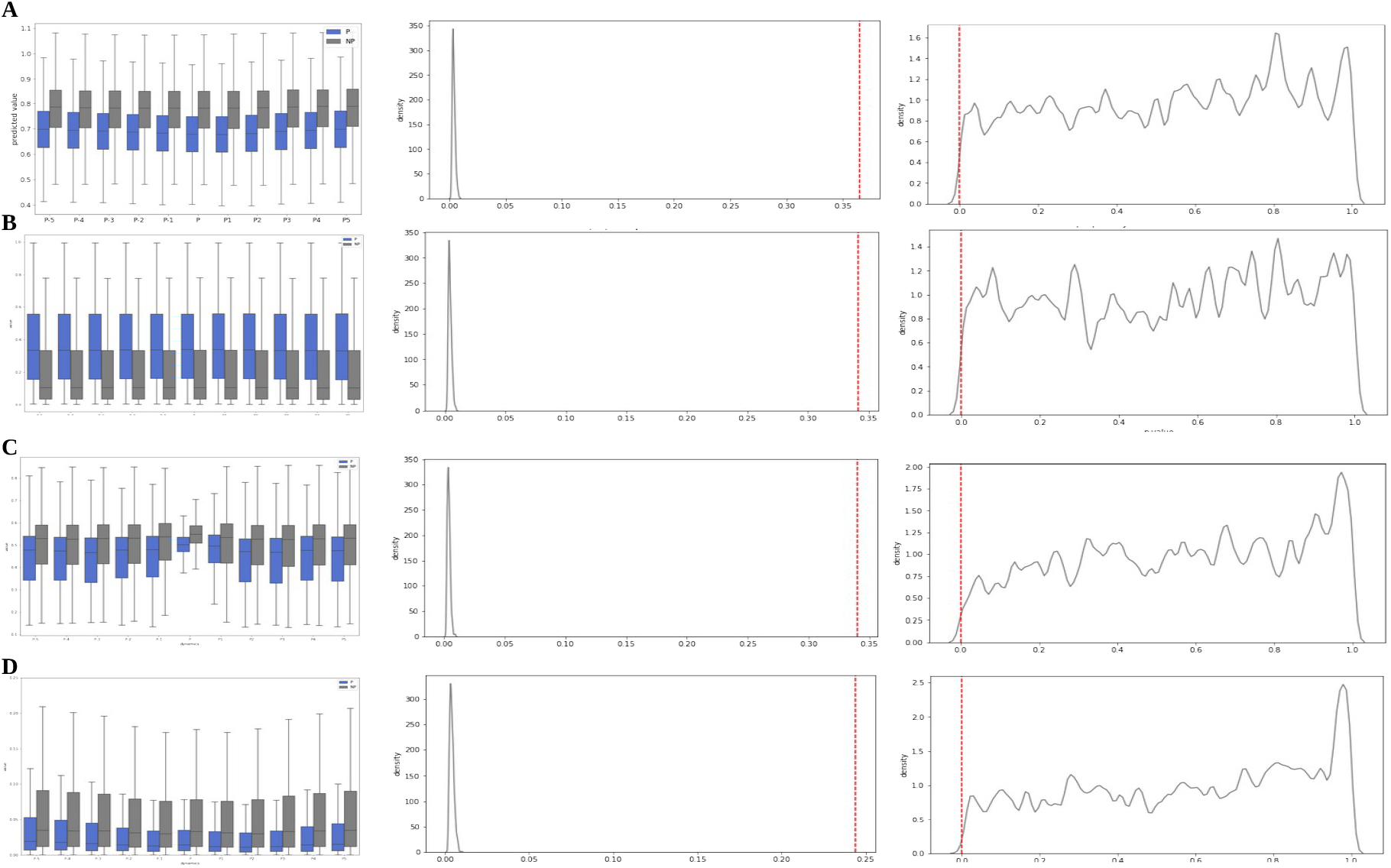
Biophysical properties of P-sites and non P-sites. **Left panel** -Distribution of dynamic values of P-sites (blue, N=79994) and non-P-sites (grey, N =1116380). **Middle panel** shows the KS-Dstat distribution of 1000 random permutation test by sampling (every permutation) same size as positive set (79994) and shuffling (red line is the actual observed value between two group) and **right panel** shows the P-value distribution of random permutation test (red line is the actual observed value) Predictions are done on primary amino acid sequence. Plot shows the biophysical preference for the sites (central site (P) and its flanking sites (P-5 to P5) that gets phosphorylated over the ones that are not phosphorylated. KS statistics are shown only for the central P-sites and non-Pistes at position ‘P’. **A)** Distribution of Backbone dynamic values. Values > 0.8: rigid, 0.69-0.8: context dependent, <0.69: flexible. **B)** Distribution of disorder propensity values. Values above 0.5indicate that this is likely a disordered residue. **C)** Sidechain dynamics: Note at ‘P’ its just S/T/Y residues and the flanking residues can be any amino acid. Higher values mean more likely rigid. These values are highly dependent on the amino acid type (i.e. a Trp will be rigid, an Asp flexible). **D)** Early folding propensity. Values above 0.169 indicate residues that are likely to start the protein folding process, based on only local interactions with other amino acids.

To check whether the above observations are real tendencies for pSTY sites, we replaced every instance of pS by A, pT by V and pY by F^70,71^ and predicted the biophysical properties for the modified protein sequence. We observed that the same sequence with modified pSTY >AVF showed an increase in rigidity; all the values of backbone, disorder, side chain dynamics and early fold propensity showed a slight increase in the modified sequence, which illustrates that it is the intrinsic nature of phosphorylatable STY residues to be more dynamic and flexible. (**Supplementary Figure S7**).

### Biophysical properties of sites in functional domains

For the biophysical analysis of P-sites and non-P-sites in functional domains, the positive and negative sets were further divided into subsets based on Pfam domain annotation, yielding four different datasets (**Figure dataset overview**).

1. P-sites in a domain region (N=20,639)
2. P-sites not in a domain region (N=30,991)
3. non-P-sites in a domain region (N=330,916)
4. non-P-sites not in a domain region (N=278,876)

To make the biophysical comparison between the subsets comparable, we only considered proteins that have sites in all four categories (7,003 proteins). **Supplementary Figure S8** shows the distributions of all biophysical properties for all four sets. The KS test was only performed for P-sites in the domain regions, and null distributions were estimated by random sampling without replacement from the subset of P-sites not in a domain region (30,991 P-sites) to match the population size of the subset with P-sites in a domain region (20,639 P-sites) (**See Methods**).

#### P-sites in functional domains tend to be more rigid

P-sites in general are more flexible than their non-phosphorylated counterparts and, as expected from the Pfam definitions, which are often structure-based, sites in domain regions are more rigid than sites that are not in a domain region (**Supplementary Figure S8:A, B**). P-sites that are in functional domains often fall in context dependent regions (DynaMine predicted value: 0.69-0.8) while P-sites that are not in domain regions are more flexible (**Supplementary Figure S8:A**). Non-P-sites that are in domains are more rigid than any other sites while non-P-sites that are not in a domain behave biophysically similar to P-sites in domain regions. The disorder propensity tends to behave similarly and in line with the backbone dynamics (**Supplementary Figure S8:B**). Side chains are more dynamic for sites that contain domain annotations. Side chains for the P-sites are less dynamic compared to non-P-sites but P-sites in functional domains have more dynamic side chains than those not in domains (**Supplementary Figure S8:C**). A similar trend was observed for early folding propensities (**Supplementary Figure S8:D**).

### Biophysical properties of P-sites with three-dimensional structural information

For the biophysical analysis of P-sites and non-P-sites with structural information, the positive and negative sets were further divided into subsets, yielding four different datasets. The sites which have at least one structure annotation in the PDB are added to set one and three and those sites without structure annotation are in sets two and four. We considered all sites that have a structural annotation (structural co-ordinates in PDB structures) of any given protein fragment length (**Figure dataset overview**).

1. P-sites with structural annotation (N=13,969)
2. P-sites with no structure annotation (N=66,025)
3. non-P-sites with structural annotation (N=164,430)
4. non-P-sites with no structure annotation (N=1,086,461)

**Supplementary Figure S9** shows the distributions of all biophysical properties for all four sets. The KS test was only performed for P-sites in the structural regions, and null distributions were estimated by random sampling without replacement from the subset of P-sites with no structure annotation (66,025 P-sites) to match the population size of the subset with P-sites with structural annotation (13,969 P-sites) (**See Methods**).

#### P-sites with structural annotations tend to be more rigid

We observed a very similar results to the P-sites with domain annotation. P-sites with structural regions are more rigid than sites that do not have structure annotation (**Supplementary Figure S9**). P-sites with structural annotations fall in context dependent regions (DynaMine predicted value: 0.69-0.8) while P-sites that does not have structural annotations are more flexible (**Supplementary Figure S9:A**). The non-P-sites that are in structural regions are more rigid than any other sites while non-P-sites that does not have structural annotation behave biophysically similar to P-sites with structure annotation. The disorder propensity tends to behave similarly and in line with the backbone dynamics (**Supplementary Figure S9:B**). Side chains are more dynamic for sites that contain structure annotations. Side chains for the P-sites are less dynamic compared to non-P-sites but the P-sites with structure co-ordinates have more dynamic side chains than those with no structure annotations (**Supplementary Figure S9:C**). A similar trend was observed on early folding propensities (**Supplementary Figure S9:D**).

### Biophysical properties of P-sites with different solvent accessibility (access, buried and interface regions)

We looked at the biophysical properties of sites with structures (subset 1 from previous analysis) that are accessible, buried, and in interface regions, to see if there are any dynamics that can explain phosphorylation in buried residues. For this analysis, we considered 6,893 P-sites that have accessibility information; 5,701 are accessible, 1,076 are interface and 116 are buried P-sites. The biophysical predictions were performed on the PDB sequences of the corresponding chains in the structures. After filtering for sites that contain five flanking residues, we retained 5,406 accessible, 990 interface, and 116 buried P-sites.

The backbone dynamics at central P-sites and flanking sites showed strong patterns for rigidity in buried sites (**Figure 12A**). The disorder propensity shows a similar pattern, but buried sites tend to be as disordered as accessible ones (**Figure 12B**). We observed an interesting pattern in sidechain dynamics and early folding propensity (**Figure 12C, D)**. The side chain dynamics of buried sites tend to be less dynamic than those of accessible and interface sites (**Figure 12C**). However, flanking sites at P-2, P-5 and P2, P4, P5 showed a similar dynamic for accessible and buried residues. The distribution of the dynamic values for accessible and buried sites become more similar as the distance from the central P-sites increases (**Supplementary Figure S10**). Even though the sidechains and backbone atoms of buried P-sites are not that dynamic, their flanking sites have very similar dynamics to accessible residues. We see a similar pattern for early folding where buried P-sites and their flanking sites from P-1 to P-4 and P1 to P4 tend to have high early folding propensity, while the distal sites at P-5, P5 have more early folding at accessible and interface sites (**Figure 12D**).

**Figure 12:**
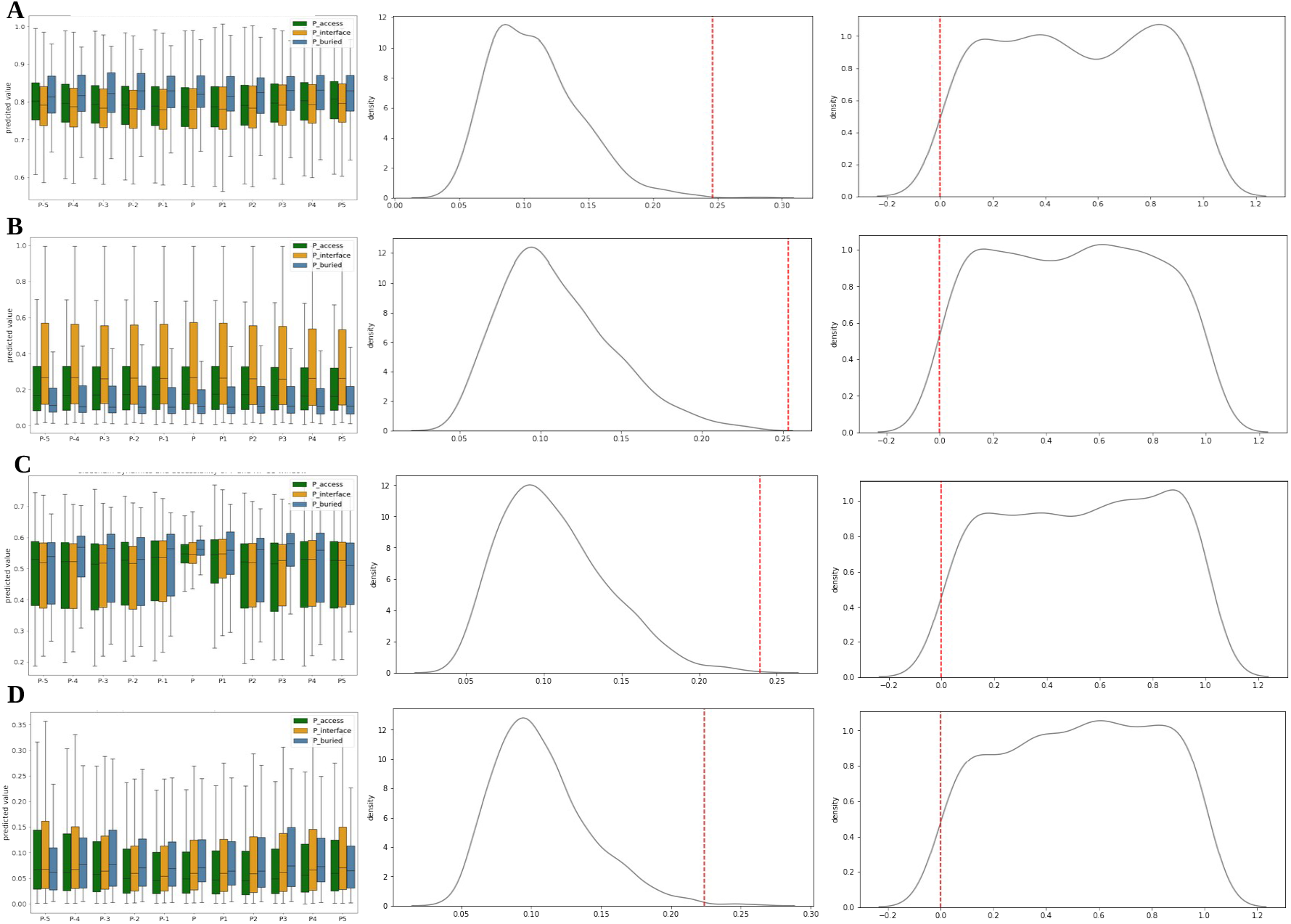
Biophysical properties of P-sites in structures from solvent exposed, interface and buried regions. **Left panel** shows the distribution of dynamic values of P-sites in solvent exposed structural regions (green, N=5406), interface regions (orange, N=990) and in buried/in-accessible regions (blue, N=116) **Middle panel** shows the KS-Dstat distribution of 1000 random permutation test by sampling (every permutation) P-sites in accessible region to match the sample size in buried regions (116) (maintained equal size for residues S:74, T:31, Y:11) and shuffling. (red line is the actual observed value between two group) and **right panel** shows the P-value distribution of random permutation test (red line is the actual observed value) Predictions are done on the correspoding PDB sequences. Plot shows the biophysical preference for P-sites. (P) and its flanking sites (P-5 to P5) in accessible, interface and buried regions of the protein structures. KS statistics are shown only for the central sites at ‘P’. **A)** Distribution of Backbone dynamic values. Values > 0.8: rigid, 0.69-0.8: context dependent, <0.69: flexible. **B)** Distribution of disorder propensity values. Values above 0.5indicate that this is likely a disordered residue. **C)** Sidechain dynamics: Note at ‘P’ its just S/T/Y residues and the flanking residues can be any amino acid. Higher values mean more likely rigid. These values are highly dependent on the amino acid type (i.e. a Trp will be rigid, an Asp flexible). **D)** Early folding propensity. Values above 0.169 indicate residues that are likely to start the protein folding process, based on only local interactions with other amino acids.

### Inter residue contacts of P-sites

Inter-residue contacts of P-sites provide information on how these sites are structurally positioned and show their degree of surface exposure in proteins. We analyzed the inter residue contacts of P-sites in the solvent accessible (5,701 sites), interface (1,076 sites), and buried (116 sites) regions by measuring the half-sphere exposure (HSE) of the residues. Half-sphere exposure (HSE) counts the number of Cα or Cβ contacts in the direction of side chains (HSE-up) and in the amino (HSE-down) direction. HSE-up values for the buried P-sites are slightly higher than the sites in accessible or interface regions (**Supplementary Figure S11**). Flanking residues around buried P-sites show varying HSE-up and HES-down values in the number of inter-residue contacts. The median contacts for the residues at P-1 and P1 show a slightly reduced HSE-up, and increased HSE-down (particularly for P1) compared to the P-site. At P-5 and P5, the median HSE-up contacts for buried residues has decreased compared to the P-site, while this has remained similar to the P-site for accessible and interface residues (**Supplementary Figure S11**). These results are in line with the biophysical properties for the central P-sites discussed above.

### Dynamic behavior of proteins and sites before and after modification

We were interested to know if there are any changes in protein dynamics in terms of residue contacts and solvent exposure before and after phosphorylation. For this, we obtained all P-sites that have evidence of phosphorylation from reprocessed MS data in Scop3P, and then restricted these to sites which had been crystallized in both states: phosphorylated (annotated as TPO, PTR, SEP residues in structure) and non-phosphorylated. For example, protein O43683 (mitotic checkpoint serine/threonine-protein kinase BUB1) has a p969S, which is crystallized in phosphorylated state in structure **6F7B: A**, and in non-phosphorylated state in structure **4R8Q: A** (**Supplementary document D2 section 1 and Supplementary Figure S13**).

We obtained 40 P-sites containing two different (phospho and non-phospho) structural states from 30 proteins and measured their HSE values. The HSE values show that the sites in non-phosphorylated states are more flexible and exposed with less residue contacts when compared to the same sites in their phosphorylated state **(Supplementary Figure S12:A).** This indicates that these sites may be more dynamic in an un-phosphorylated state and upon phosphorylation they form more contacts making it less flexible. The HSE contacts of flanking residues show an alternate pattern, for example the immediate flanking sites P-1 and P1 have more residue contacts towards the backbone (HSE-down) and less side chain contacts (HSE-up) after modification (**Supplementary Figure S12:Bii, iii))** while an opposite trend was observed for P-2 and P2 sites (**Supplementary Figure S12: Bi, iv)**). This alternating backbone and side chain flexibility in the flanking residues for unmodified P-sites can favor phosphorylation of the P-site. This because the compound flexibility of the flanking residues can expose an otherwise buried P-site, which, upon phosphorylation, will form more contacts, thus becoming more stable.

#### Possible dynamic behavior of P-sites

Combining all these observations, the possible dynamic nature of buried P-sites are explained in the supplementary information **(Supplementary document D2 section 2)** for three proteins; beta actin (ACTB) (**Supplementary Figure S14)**, Phosphoglycerate kinase 1 (PGK1) (**Supplementary Figure S15)** and Glyceraldehyde-3-phosphate dehydrogenase (GAPDH) (**Supplementary Figure S16)**.

## Discussion

A better understanding of post-translational modifications (PTMs) of proteins is essential to improve our knowledge of the signaling networks in cellular mechanisms, with protein phosphorylation particularly relevant. Mass spectrometry based proteomics studies have revolutionized the identification of large numbers of such phosphorylation sites in proteins through large scale proteome wide experiments. However, the large amount of data that are available are of varying quality and reliability, as they have been typically generated and analyzed over many years, using different algorithms and approaches. As a result, P-sites from publicly available data may not have been localized, may have been identified against outdated protein sequence databases, or may have been identified using a sub-optimal search engine. In this work, we therefore used consistent state-of-the-art re-processing of the original raw data from public proteomics data to obtain more consistent and robust results for protein phosphorylation sites, with reliable P-sites in human proteins distinguished from random sites based on combinatorial information obtained from different resources and associated meta-data. After filtering for multiple evidence and proteomics derived meta-information for individual P-sites, we so obtained 81,404 sites as reliable P-sites, which together cover 60% of canonical human proteins. From our analysis, we found that phosphoproteins show a distinct pattern based on the phosphorylated residue types (**Figure 5C-D**); some proteins are phosphorylated on specific amino acid types (pS/pT/pY), while others are phosphorylated on all residues (pSTY), with the latter tending to be more abundant. Interestingly, we observed that the relative phosphorylation of Tyr is as frequent as Thr phosphorylation in our data.

Our analysis enables a more reliable view on the proteome-wide P-sites found in human samples, and we further placed this information in the context of the sequence, structure, and biophysical properties of proteins, to revisit previous analyses in relation to phosphorylation. Several studies have previously reported that phosphorylation occurs more often in the protein termini^16,72^ due to the typically higher conformational flexibility; our analysis here shows that P-sites are in fact spread out evenly along the protein sequence, with only a slight increase observed in the C-terminal regions (**Figure 6**). C-terminal regions of proteins often contain short linear motifs (SliMs), and are modified by several PTMs, especially phosphorylation, so mediating protein-protein interactions and sub-cellular localization. This conformational flexibility and their importance in cellular regulations might cause the C-termini regions to be more prone to phosphorylations, and possibly other PTMs.

Though we observe more than 60% of the human proteome to undergo phosphorylation, only 5% of sites are annotated to have functional relevance^40,45^, often using protein domains and structural information^42,73^ of P-sites and proteins. Our analysis of these P-sites in their protein-domain context shows that, although the most widely represented domain families in the proteome were most often found in our data, domains with less frequent occurrences in proteins tend to have more phosphorylated sites. With 10% of all STY sites in the proteome observed to be phosphorylated, only 2% of them are in domain regions, and the remaining 8% falls in the inter-domain regions of the protein. Most of these inter-domain regions are unstructured and flexible, and are likely to experience conformational switches like folding upon binding. It is therefore important to know their structural and biophysical characteristics to understand their regulatory roles upon phosphorylation.

Proteins are indeed dynamic in nature, which makes the physicochemical and biophysical consequences of phosphorylation challenging to study. Studies have shown that phosphorylation can induce local and global changes in protein structures^74^ and often undergo structural transitions (order ↔ disorder) upon phosphorylation or de-phosphorslation events^25,26^. Recently ***Henriques J and Lindorff-Larsen K***^75^ showed how an inaccessible Tyr residue is exposed upon phosphorylation through molecular dynamics simulation analysis. We considered the structural, non-structural (disorderedness) and biophysical context of the proteins and P-sites to obtain more insight on the structure-function relationship of proteins and the possible conformational changes. Our data suggests that most of the P-sites are in flexible, unstructured regions, with only 40% of P-sites observed in regions with helical or beta-strand propensities. It has been previously reported that phosphorylation occurs in solvent accessible, flexible regions of proteins^16,20,21^. Though most of the P-sites are in solvent accessible regions, we found a considerable number of sites (15% of all P-sites with accessibility information) that are in interface or in solvent inaccessible/buried regions (**Figure 8A**).

Our analysis of the biophysical properties of these sites and their flanking regions shows that these buried residues are typically surrounded by solvent accessible residues that have flexible and dynamic properties, potentially allowing increased accessibility of these P-sites through the resulting dynamics. Moreover, sites that are buried in their un-phosphorylated state tend to be in the regions that have structural transition properties like disorder↔order, molten globule↔globule and folding upon binding. These sites are interesting as they typically are in a ‘context-dependent’ state of dynamics with the possibility to be either rigid or flexible based on their environment. Our data therefore offers new opportunities to understand where conformational changes upon modifications and/or mutations might take place. Combining the physical residue contacts from HSE (decrease in contacts with increase in flanking distance) around buried sites (**Supplementary Figure S11**) and predicted biophysical properties (backbone and side chain dynamics (**Figure 11, Figure 12**), we conclude that buried yet phosphorylated P-sites are typically sufficiently dynamic to undergo conformational changes that expose these sites.

We confirmed the functional relevance of P-sites are through a analysis of observed mutations. Such mutation of P-sites may disrupt the protein structure, causing loss or gain of functions and introducing potential new P-sites. Since most of the P-sites are in unstructured regions of the proteins, it is possible, but less likely that a phospho mutant might cause a deleterious effect. Our analysis on mutational hotspots shows that variations on and around P-sites are more neutral when compared to variations on non-P-sites. We also see that mutations occurring on rigid P-sites (minimal dynamics) more often cause deleterious effects upon mutation.

In summary, our analysis is the first and largest scale study with an integrated approach to understand the P-sites and phosphoproteins by placing them in a variety of possible contexts such as sequence, structure, disorderedness and biophysical properties. With the availability of large amounts of proteomics data and the increase in identification of protein PTMs, we show how re-processing of available proteomics data can help prioritize relevant P-sites from random ones based on combinatorial meta-data evidence. Though our analysis was focused towards protein phosphorylation, we envision that this can be extended to other modification types like ubiquitination, acteylation and methylation. This will help the scientific community understand co-translational modifications and PTM cross-talk in cellular regulation. As more re-processed proteomics data and associated meta-information like sub-cellular localization becomes available, it will further help pinpointing the stoichiometry of modifications inside the cell. Proper validation on the dynamic behavior of protein structures through nuclear magnetic resonance (NMR) experiments or by molecular dynamics simulation (MD) studies on highly relevant P-sites might help to unravel interesting dynamic behavior of residues modified in the solvent inaccessible regions. While the biophysical and structural characterization of phosphorylation events is still in its early stages, from the observed data it is clear that such endeavors are worthwhile, especially given phosphorylation’s key importance in signal transduction, protein binding, and function.

## Conflict of interest

The authors declare that they have no conflict of interest.

## Acknowledgments

PR, LM, WV acknowledge funding from the Research Foundation Flanders under grant agreement number G.0328.16N. LM acknowledges funding from the European Union’s Horizon 2020 Programme under Grant Agreement 823839 (H2020-INFRAIA-2018-1).

## Supporting information

**Supplementary Table T1**

All filtered confident P-sites (positive set) and the meta-data used in the analysis.

**Supplementary Table T2**

Functional domain annotation of P-sites for every protein instance

**Supplementary Table T3**

Average phosphorylation status for every domain by pooling the protein instances together.

**Supplementary document D1**

Contains all the supplementary figures

**Supplementary document D2**

Case studies and figures showing the dynamic nature of P-sites and proteins

## Notes

https://github.com/Pathmanaban/Phospho_analysis

### Competing Interest Statement

The authors have declared no competing interest.

