## Supplementary Document D2 for "A panoramic perspective on human phosphosites"

### 1. Dynamic behavior of proteins and sites before and after modification in BUB1A protein

We investigated the phosphorylated and non-phosphorylated protein structure of protein **BUB1A**: a mitotic checkpoint serine/threonine-protein kinase, which has 29 P-sites in our data. The site p969S is mapped to the protein structure (6F7B: A) which has been crystallized in phosphorylated state at the same position in structure (SEP 969). We obtained the protein structure (4R8Q: A) for the same protein but in a non-phosphorylated state at the same position (SER 969). Our HSE analysis shows that in the non-phosphorylated state (4R8Q) SER 969 makes six inter-residue contacts in the direction of the side chain (HSE up) and 14 contacts in the direction of the amino backbone (HSE down). In phosphorylated state (6F7B) site p969S/SEP 969 has 11 contacts in the side chain direction (HSE up) and maintains the same 14 contacts in the backbone direction (HSE down). The superposed structures show a clear distinction of the central site and its flanking regions (phosphorylated 6F7B in pale green and un-phosphorylated 4RQ8 in pale blue) (**Figure S13**). In phosphorylated state, the central P-site p969S (red sticks) and the flanking regions (pale green) face more towards the structural core allowing more interactions. The side chain of p969S extends close towards the neighboring amino acids in structural space, while the un-phosphorylated Ser 969 (blue sticks) and the flanking stretch are facing away from core, favoring towards the direction of solvent accessible region (**Figure S13**). This shows that Ser 969 in its un-phosphorylated form is more solvent exposed that might allow kinase to phosphorylate and once phosphorylated the same site makes additional contacts making it less exposed, indicating the dynamic behavior of the P-site before and after phosphorylation.

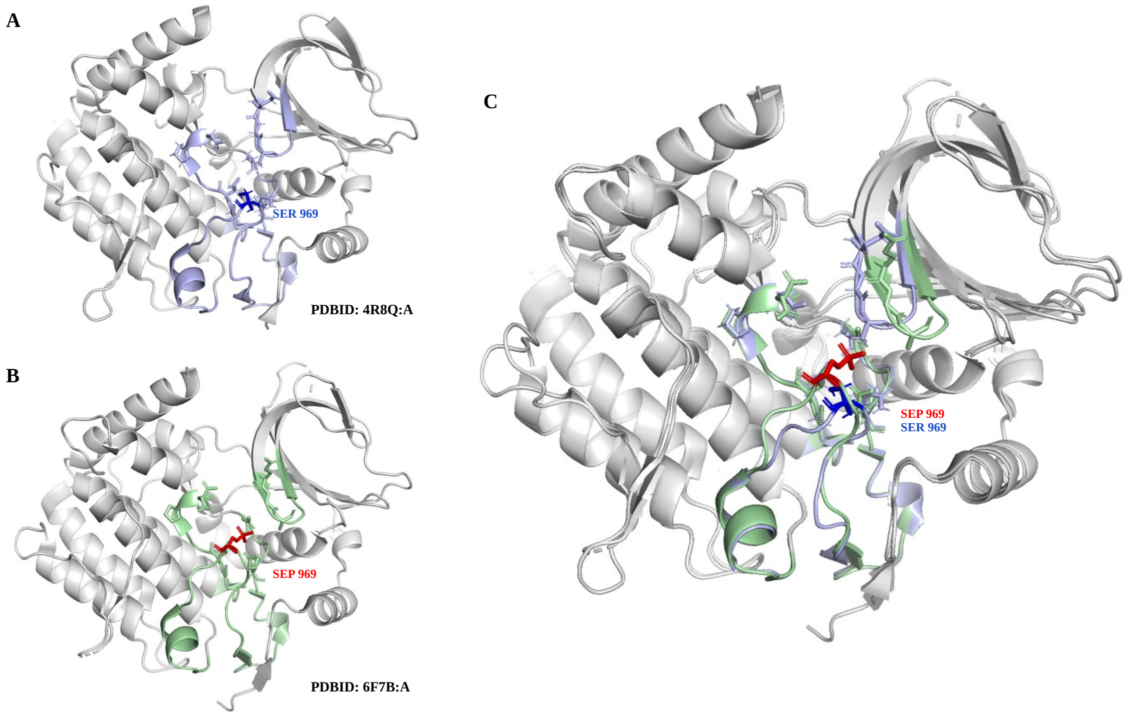


**Figure S13: mitotic checkpoint serine/threonine-protein kinase BUB1 protein in phosphorylated and un-phosphorylated form at SER 969**
P-site p969S from O43683 protein mapped to two structures: 4R8Q:A where Ser 969 is in un-phosphorylated state and 6F7B:A where Ser 969 is in phosphorylated state.

**A)** Structure with un-phopshorylated Ser at position 969 (SER 969). The central site Ser 969 and its sequence and structurally close residues are colored in blue and shown in stick representation **B)** structure with phosphorylated Ser at position 969 (SEP 969). The central site SEP 969 and its sequence and structurally close residues are colored in red and shown in stick representation. **C)** Superimposed structures showing the extended side chain of SEP 969 (red) towards the spatial neighbors and the flanking amino acids favoring the interaction and un-phosphorylated SER 969 an flanking sites favoring less contacts and more exposed.

### 2. Combining Structure, biophysical properties and structural transition annotations to explain the dynamic nature of buried P-sites

We looked at three different proteins and the identified P-sites combining all the structural and biophysical information in our data.

#### i) **Beta actin/ACTB (P60709)**

Beta actin is a structural protein that is involved in key cellular functions like contraction and motility. This protein is highly phosphorylated in our data having 33 P-sites, which makes that 50% of all STY sites in this protein can be phosphorylated. All these sites are mapped onto the protein structure 3BYH: A (**Figure S14A**). Only one solvent inaccessible site was seen for this protein which is p32S (in the structure this residue has position 33) (indicated by dashed blue line/circle) (**Figure S14A, B**). The site shows rigid backbone dynamics but flexible side chains, moreover our structural transition annotation from MobiDB shows that this site and its flanking regions (position 33-78) have disorder to order transition properties (shown in cyan in **Figure S14A**) indicating that, upon binding with a kinase in the process of multi phosphorylation or with other proteins during protein-protein interaction, this site might get exposed for phosphorylation.

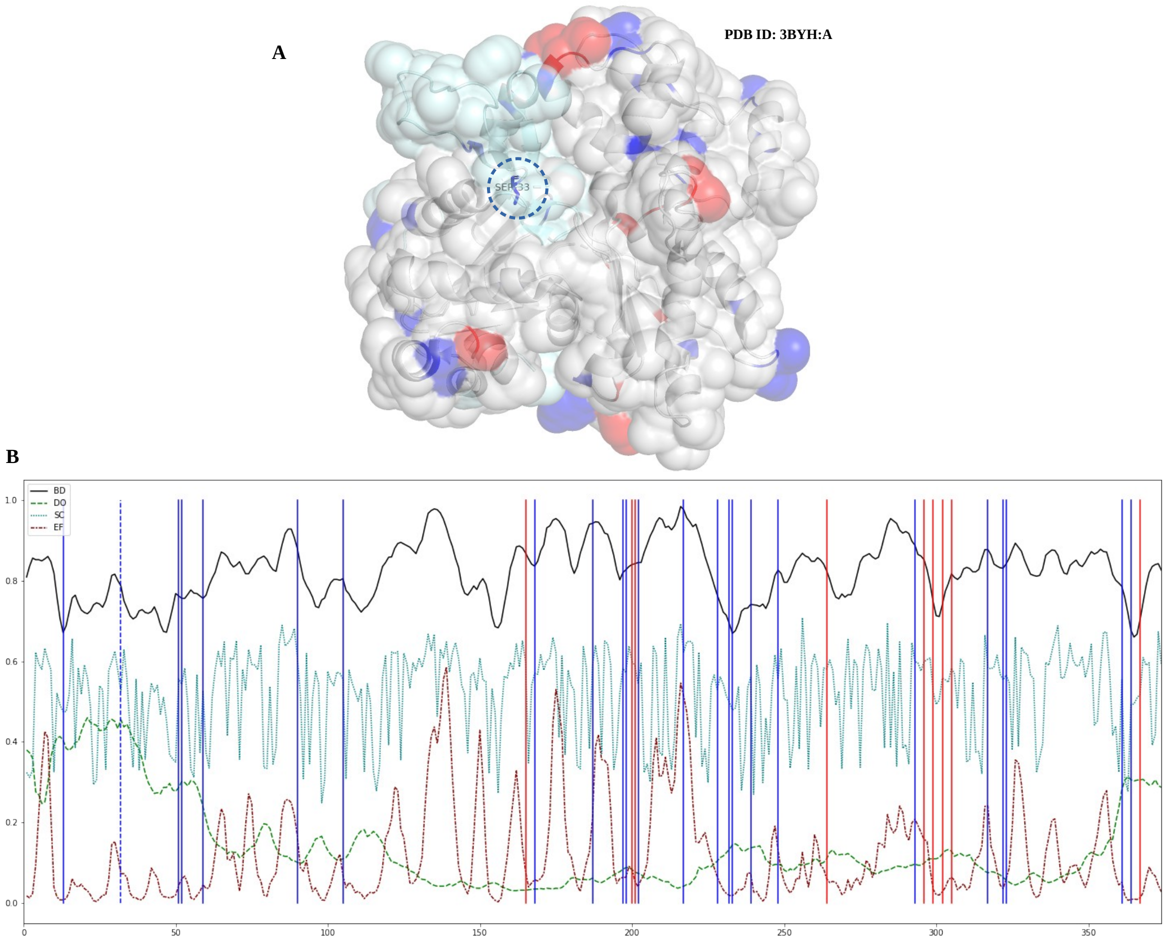


**Figure S14: Structure and dynamics of Beta-actin**
**A)** P-sites with structural co-ordinates for protein P60709 mapped onto the protein structure 3BYH:A. MPS sites are colored blue and SPS are colored red. Protein structure is rendered with solvent accessible surface area and in cartoon representation with 60% transparency. Buried residues is indicated in dotted blue circle in structure. Pale cyan in structure indicates the region is an intrinsically disordered region (IDR) or evidence for any structural transition (order-disorder, disorder-order) observed upon protein protein interaction

B) The biophysical predicted values are shown. Backbone dynamics (BD) colored in black, disorder propensity (DO) colored in green, side chain dynamics (SC) colored in cyan and early folding propensity (EF) colored in maroon. As in structure the MPS are colored blue and SPS in red. Buried sites are indicated with dashed lines.

ii) **Phosphoglycerate kinase 1/PGK1 (P00558)**

Phosphoglycerate kinase plays a key role in the glycolytic pathway, as a primer recognition protein, and plays a role in sperm motility. This protein contains 10 P-sites in our data and all these sites are mapped onto the protein structure 2WZB: A. Three of these sites (57S, 62S, and 175S in sequence, corresponding to p56S, p61S and p174S in structure) are in the solvent inaccessible region of the protein (**Figure S15A, B** indicated by dashed blue line/circle). There is no known annotation of IDRs or structural transition annotations for this protein. Structural annotation and the biophysical values show that the site p61S is situated in a flexible loop (**Figure S15A**) where the region N-terminal to the site is more flexible (flexible dynamic values) and adopts a loop conformation (in structure) (**Figure S15B**). Also, p61S is structurally close to the other solvent inaccessible site p174S (**Figure S15A**). The secondary structure of p174S adopts a helical conformation and is seen in a coil-helix-coil form (helix formation for residues 173-176), the biophysical values show alternating side chain dynamics in the flanking residues around the p174S site. Site p56S is located structurally close to p114S, and the N-terminal flanking regions of this residue tends to be more flexible (adopting a loop conformation in structure (**Figure S15A**)) with flexible dynamic values (**Figure S15B**), indicating that these dynamics are involved to expose such buried sites to phosphorylation upon multi-site phosphorylation events.

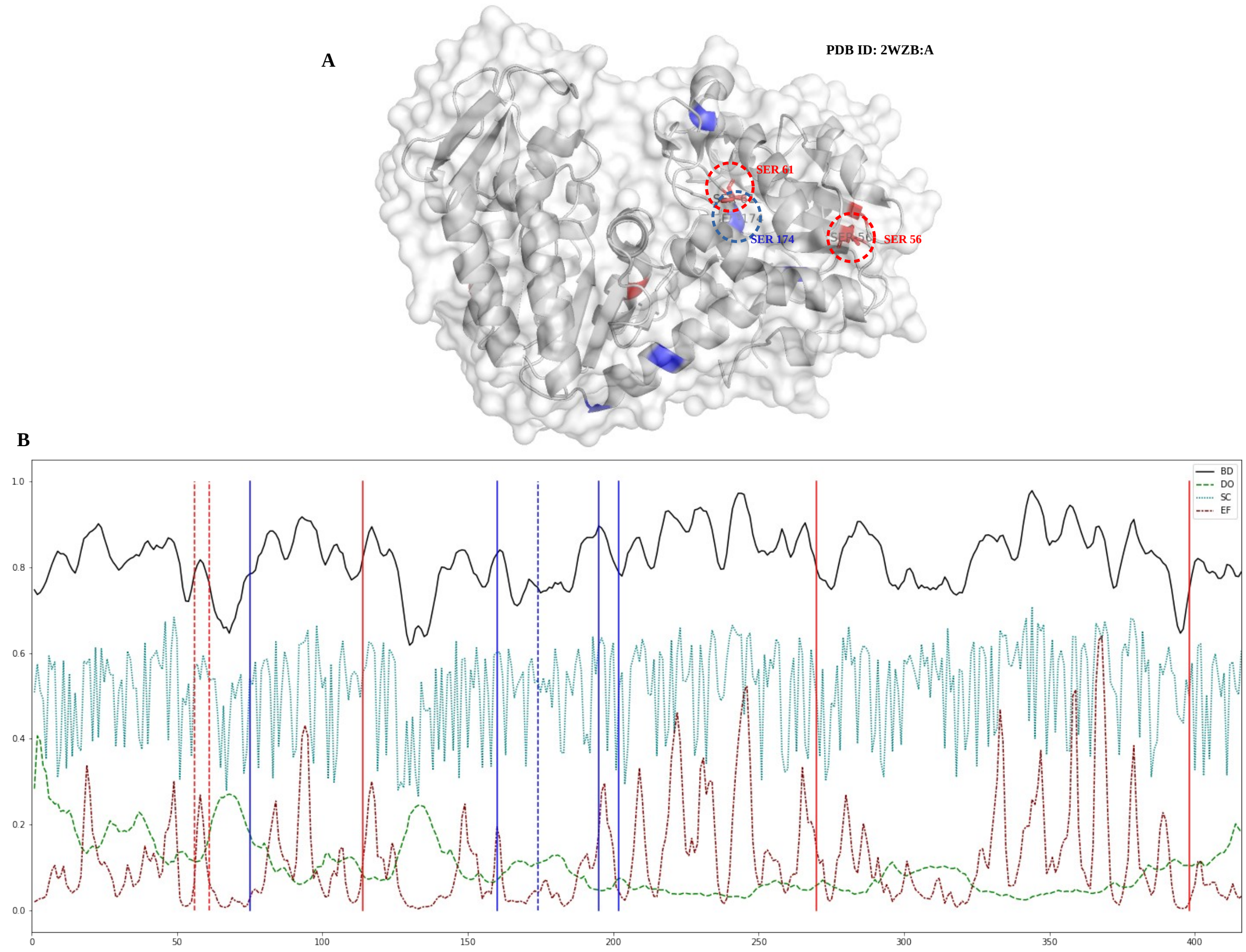


**Figure S15: Structure and dynamics of Phosphoglycerate kinase 1**
**A)** P-sites with structural co-ordinates for protein P00558 mapped onto the protein structure 2WZB:A. MPS sites are colored blue and SPS are colored red. Protein structure is rendered with solvent accessible surface area and in cartoon representation with 60% transparency. Buried residues is indicated in dotted blue circle in structure. Pale cyan in structure indicates the region is an intrinsically disordered region (IDR) or evidence for any structural transition (order-disorder, disorder-order) observed upon protein protein interaction

B) The biophysical predicted values are shown. Backbone dynamics (BD) colored in black, disorder propensity (DO) colored in green, side chain dynamics (SC) colored in cyan and early folding propensity (EF) colored in maroon. As in structure the MPS are colored blue and SPS in red. Buried sites are indicated with dashed lines.

iii) **Glyceraldehyde-3-phosphate dehydrogenase/GAPDH**

GAPDH is a key protein in the cell, as it is involved in various cellular processes and regulation including transcription, RNA transport, DNA replication and apoptosis. Our data contains 21 P-sites for this protein, which makes that 40% of all STY sites in this protein can be phosphorylated. All these P-sites are mapped onto the protein structure 6YND: A (**Figure S16A**). Site p148S is seen in the solvent inaccessible region of the protein structure (**Figure S16A** indicated by dashed blue line/circle). This site has a beta strand secondary structural element formed by three residues (from 145-148) and the N and C terminal regions to the site form flexible loops. Moreover, the site is hindered by a large flexible loop (from 122-136) which forms an arc like formation over p148S (**Figure S16A**). The biophysical values show that N-terminal regions of the immediate flanking residues and the sequentially farther but structurally close (arc like loop forming residues in structure) residues are flexible in both backbone and side chains, favoring higher disorder propensity (**Figure S16B**). Site p148S is itself anchored between two other P-sites p143S and p151S, and the P-site p125S in the arc like loop (**Figure S16A, B**). The two anchoring sites (p143S and p151S) are in flexible loops. There are some IDRs observed for this protein but they are both sequentially and structurally far from these sites (**Figure S16A** indicated by pale cyan). Looking at the consecutive Ser phosphorylation around the buried p148S and the flanking flexible regions around this site, we assume there might be a series of phosphorylation events by a Ser/Thr kinase that lead to exposure of the p148S site for phosphorylation.

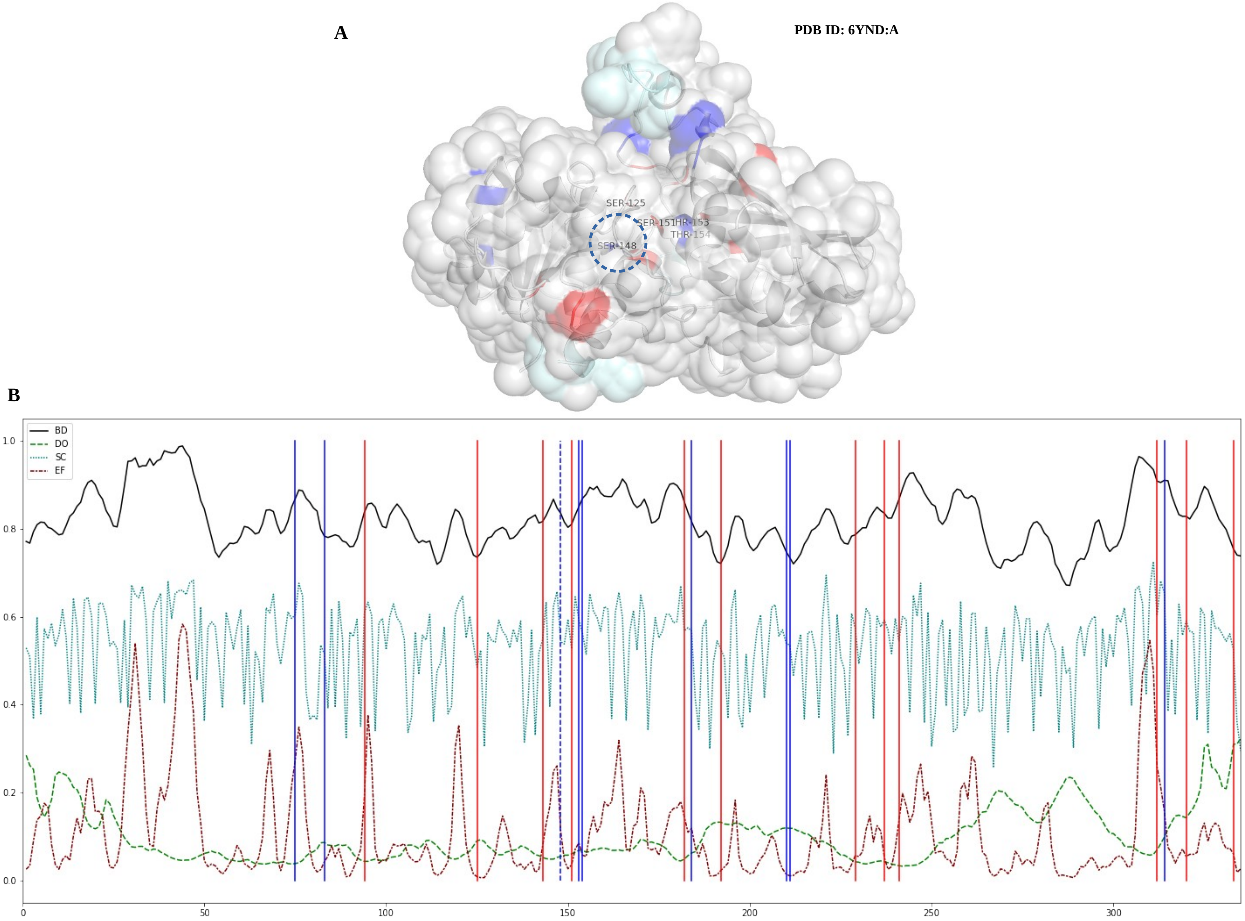


**Figure S16: Structure and dynamics of Glyceraldehyde-3-phosphate dehydrogenase**
**A)** P-sites with structural co-ordinates for protein P04406 mapped onto the protein structure 6YND:A. MPS sites are colored blue and SPS are colored red. Protein structure is rendered with solvent accessible surface area and in cartoon representation with 60% transparency. Buried residues is indicated in dotted blue circle in structure. Pale cyan in structure indicates the region is an intrinsically disordered region (IDR) or evidence for any structural transition (order-disorder, disorder-order) observed upon protein protein interaction

B) The biophysical predicted values are shown. Backbone dynamics (BD) colored in black, disorder propensity (DO) colored in green, side chain dynamics (SC) colored in cyan and early folding propensity (EF) colored in maroon. As in structure the MPS are colored blue and SPS in red. Buried sites are indicated with dashed lines.
