## Supplementary Document D1 for "A panoramic perspective on human phosphosites"

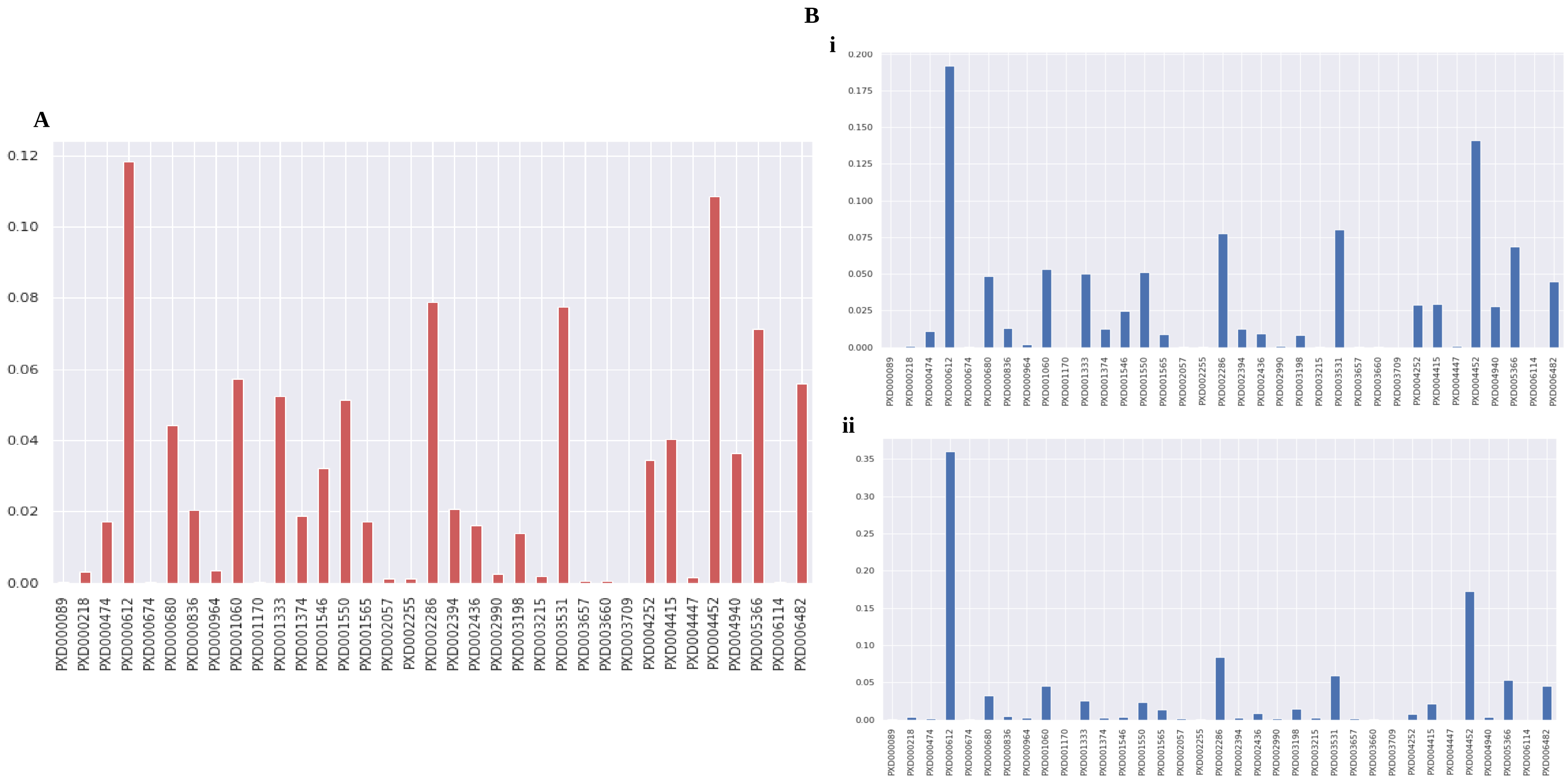


**Figure S1: Properties of the SPS**

**A) P**roject frequency of the proteins containing SPS. Density of total number of proteins (y-axis) with the given project frequency (x-axis). There are some proteins seen just in one project but most of the proteins are seen in two or more projects. **B)** Total number of P-sites identified per project in Scop3P **(i)** to the total number of SPS P-sites seen in the same project **(ii)** PRIDE project IDs in x-axis and log transformed number of P-sites in y-axis

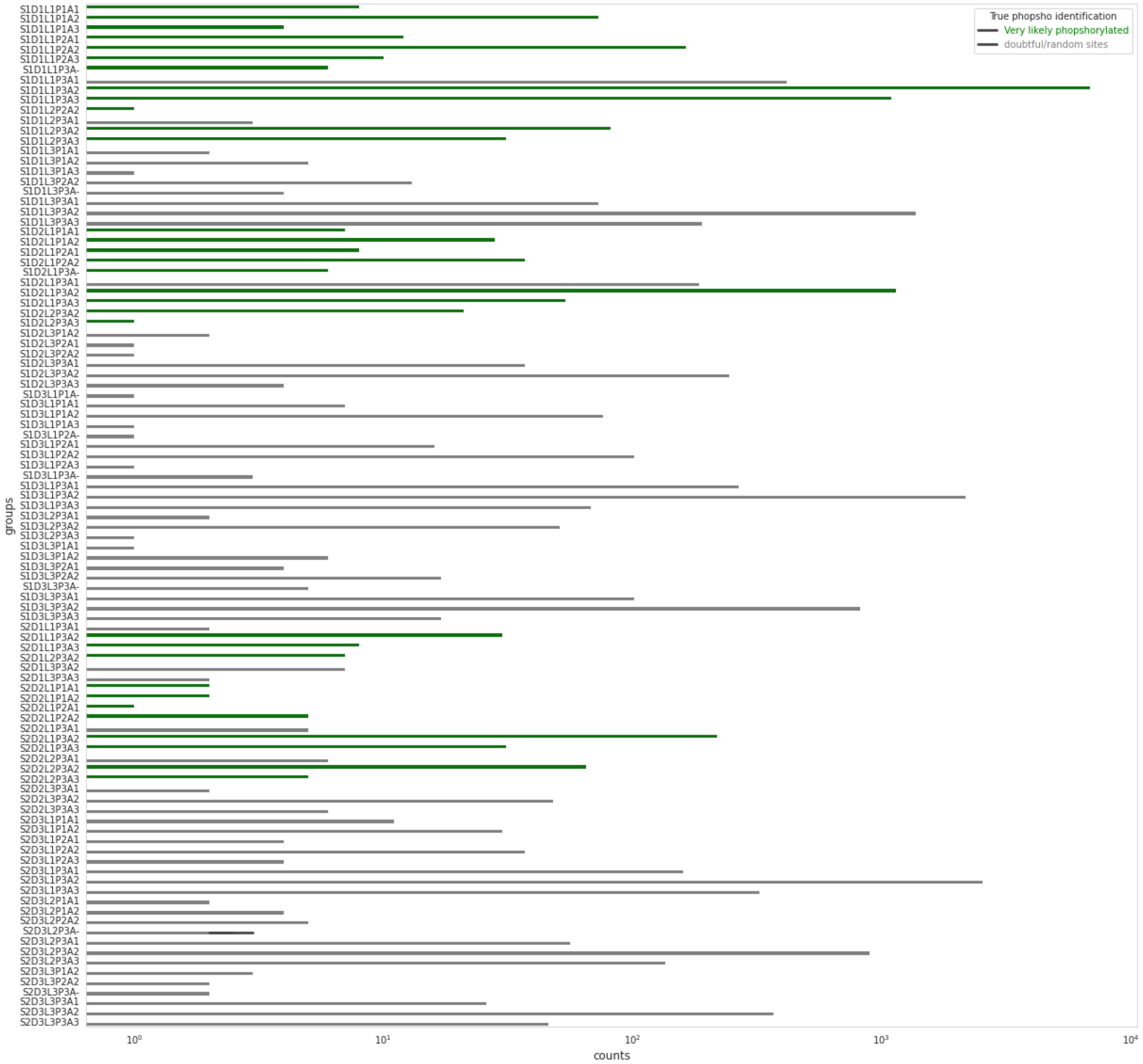


**Figure S2: Filtering reliable P-sites from random ones:**

Number of reliable P-sites (green) and random ones (grey) are shown in x-axis and y-axis the groups used to filter them based on five different site meta-data: protein abundance, phospho spectra frequency, distance to the nearest P-sites identified, site localization probability and phosphorylation status on the identified peptide (single or multi). For example a site seen in multi phosphorylated peptide (S2) with distance of >20 amino acid apart (D1) having localization probability of >70% (L1) with the <5 spectral frequency (P3) but the protein is not abundant (A3), we consider the sites as true phosphorylation (S2D1L1P3A3) but for the same site with all same criteria but if the protein is high abundant (S2D1L1P3A1) we consider that these sites are possibly random occurrence based on the fewer spectra from a highly abundant protein. Based on such combinations the sites colored green are considered reliable P-sites and the grey ones are possible noises in the data. x-axis shows the log transformed P-sites counts in each category in y axis.

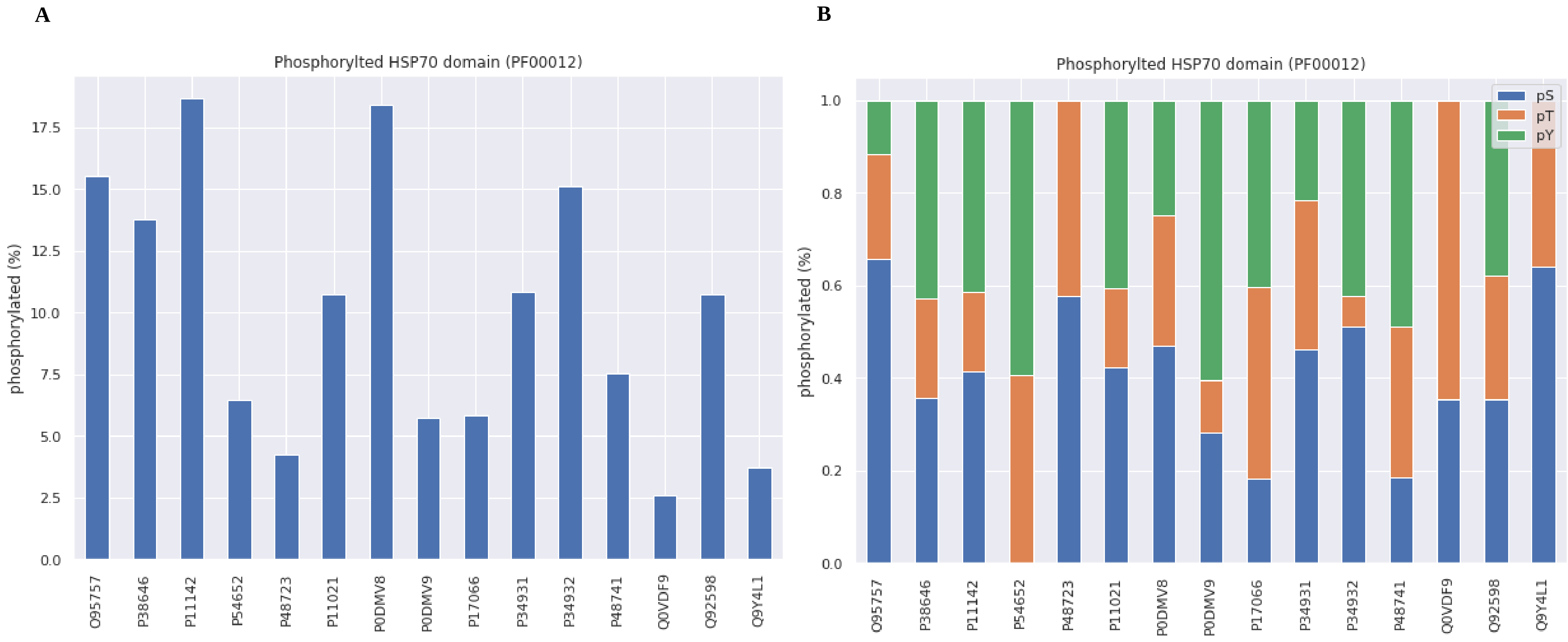


**Figure S3: Phosphorylation on HSP70 domain**

The difference in phosphorylation seen in different protein instances (x-axis) of the same domain (HSP70) because of difference in sequence composition in proteins **(A)** and the type of amino acid phosphorylated in different instance **(B)** Different protein instance (UniProt protein ID) of the domain is shown is x-axis and the percentage of phosphorylation on y-axis

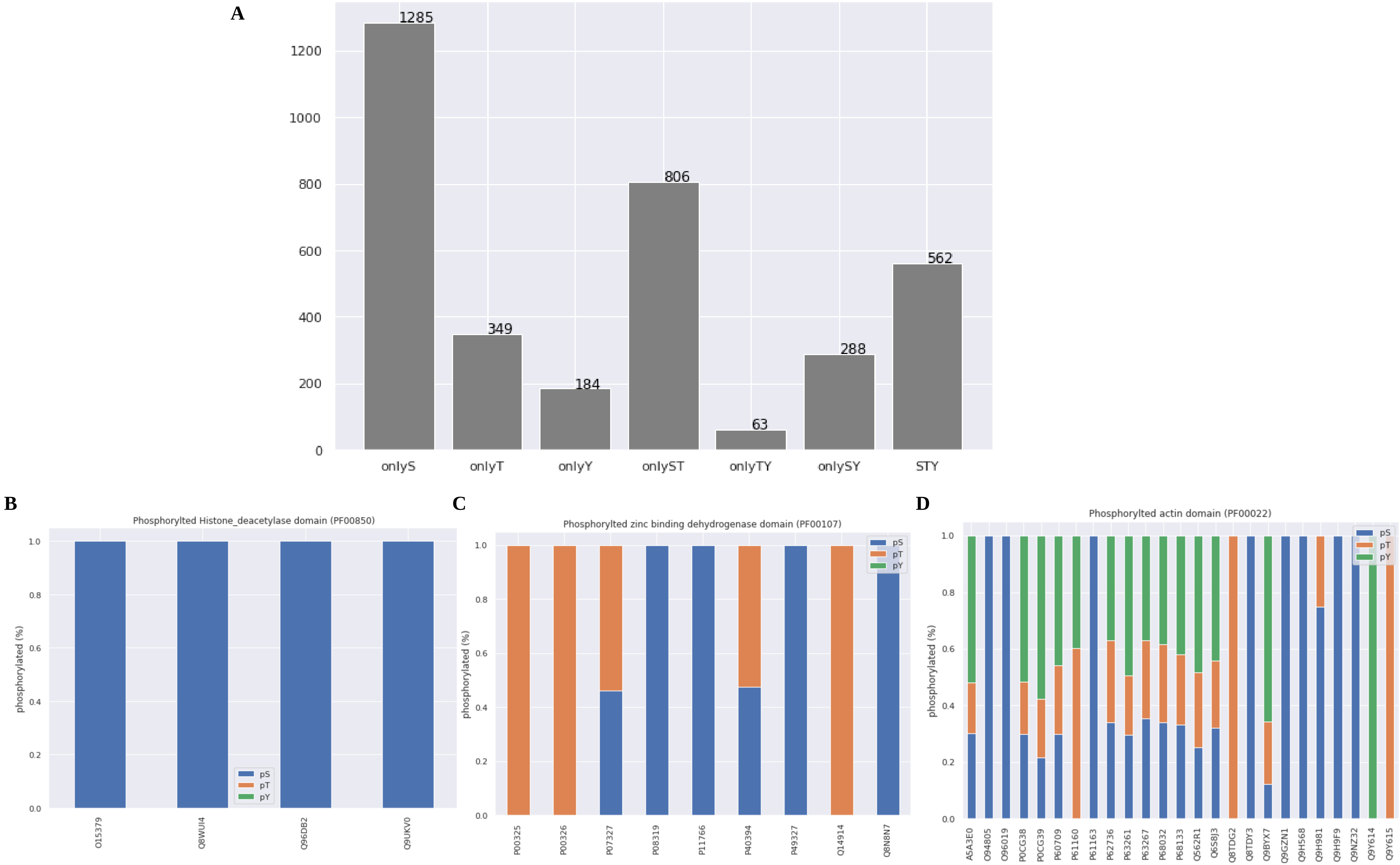


**Figure S4: Residue specific domain phosphorylation**

**A)** The breakdown of number of domains (y-axis) seen as phosphorylated in specific residues (S,T,Y), in combination of different residues (ST, SY, TY) or in all STY residues **B,C,D)** Examples of domains that are seen as phosphorylated in all instance at Ser residue only (**B**-PF00850: histone deacytilase domain), and phosphorylated in Ser/Thr in all or any instance (**C** -PF00107: zinc-binding dehydrogenase) and seen as phosphorylated in all STY in any or all instance of the domains (**D**-PF00022: actin domain). Different protein instance (UniProt protein ID) of the domain is shown is x-axis and the percentage of phosphorylation in y-axis.

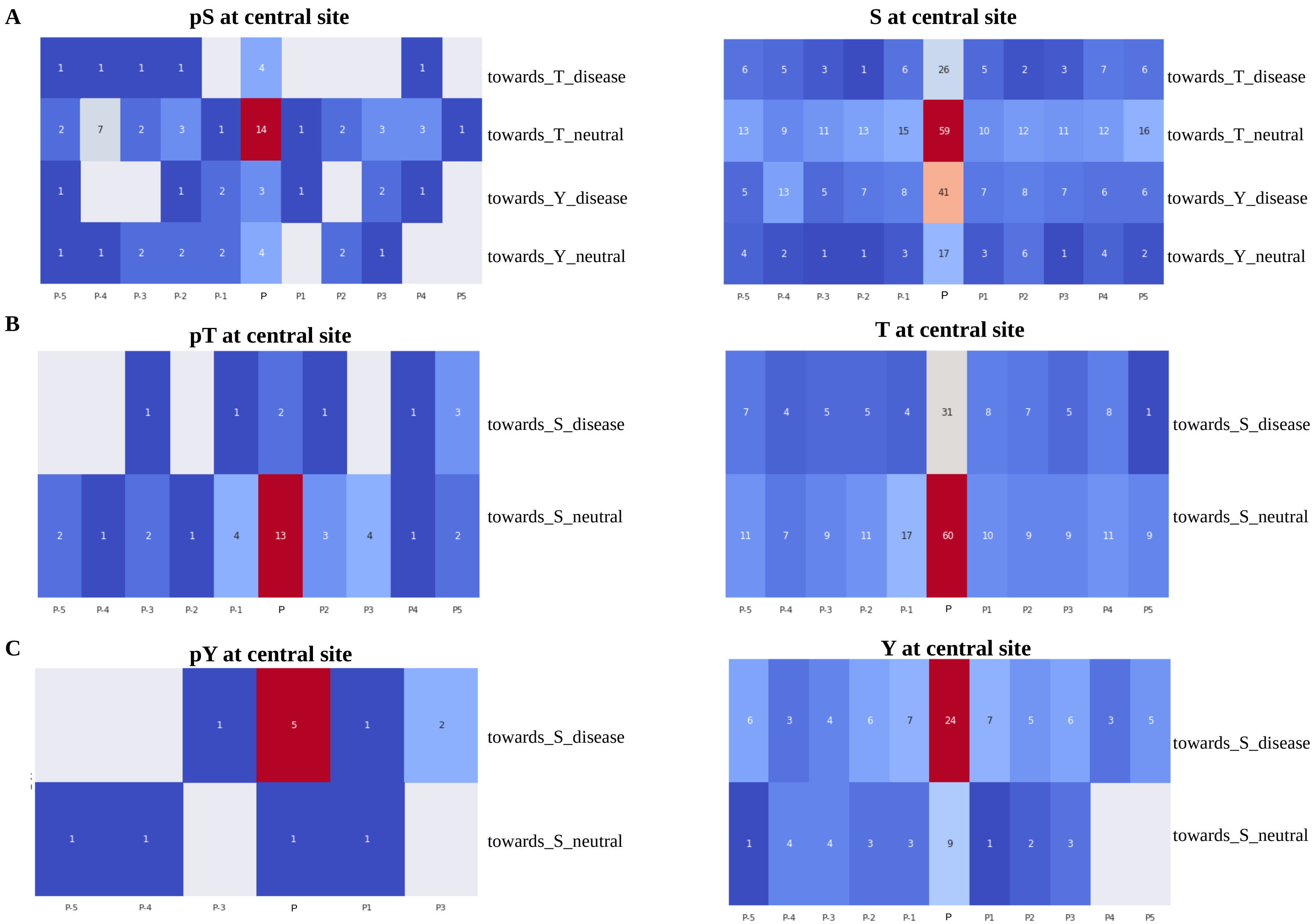


**Figure S5: Self phospho variants on and around P-sites and non P-sites**
Illustartes the frequency of amino acid variants in central (P) and its flanking N (P-5 to P-1) and C terminal (P1-P5) sites around P-site (left panel) and non P-sites (right panel). The color range from blue (less frequency) to red (more frequency). X axis shows the location of mutation. P denotes the mutation is seen is central site (**A: pS/non-phospho S, B: pT/non-phospho T and C: pY/non-phospho Y**) and P1 means there is a mutation next to a P-site in the C terminal position and P-5 denotes a mutation at $5^{th}$ residue in the N-termini when there is P-site at position 'P'. Y axis shows the variant amino acid type for every position followed by if the variant is disease causing or neutral. Every cell in the plot represents the frequency of mutation seen at that position.

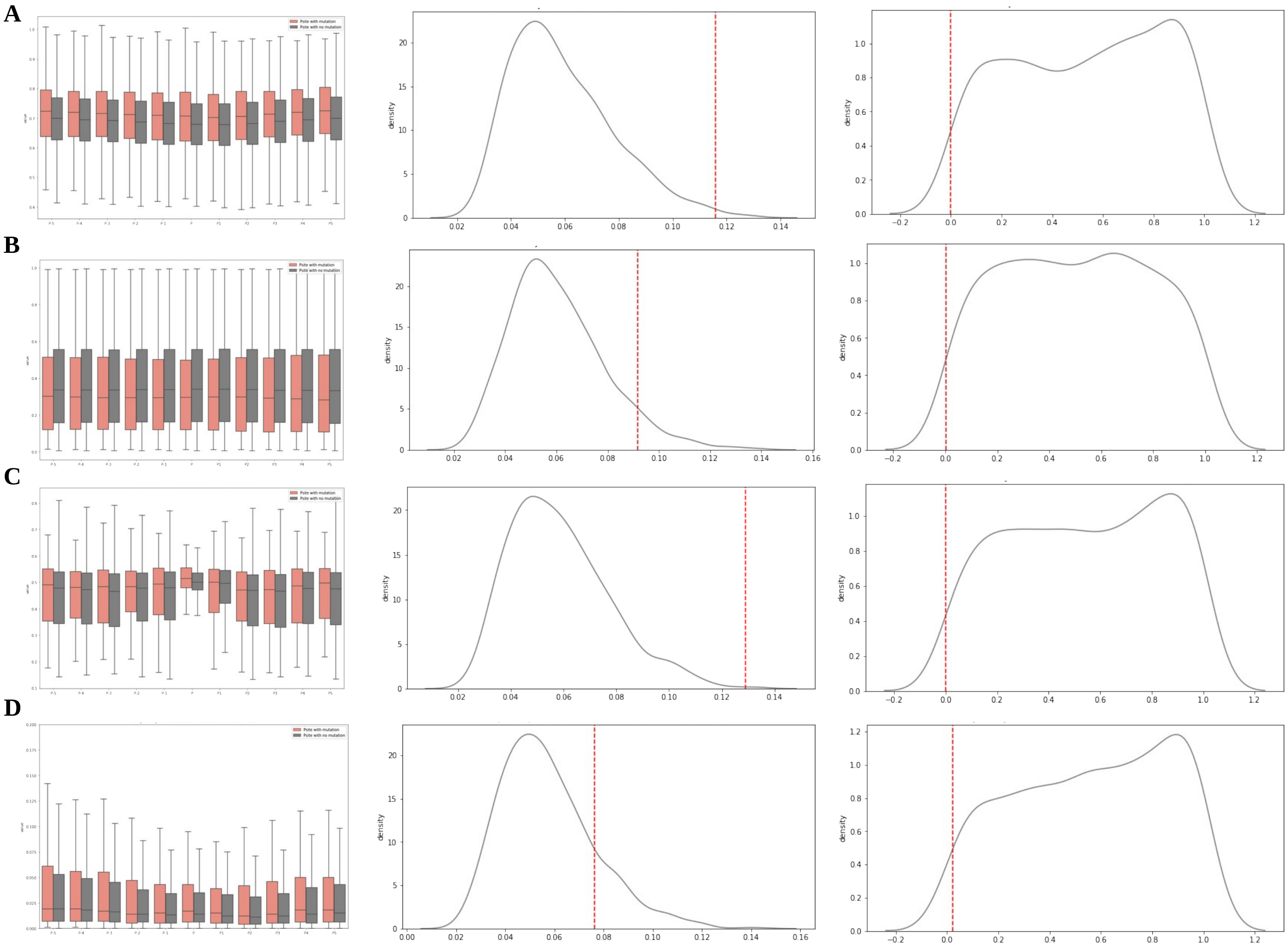


**Figure S6: Biophysical properties of mutation at P-site vs mutations not in P-sites**

**Left panel** -Distribution of dynamic values of variants at P-sites (Salmon, N=389) and variants not in P-sites (dark grey, N=79606). **Middle panel** shows the KS-Dstat distribution of 1000 random permutation test by random sampling (every permutation) of variants not in P-sites to match the sample size of phospho variants (N=389) and shuffling (red line is the actual observed value between two group) and **right panel** shows the P-value distribution of random permutation test (red line is the actual observed value) Predictions are done on primary amino acid sequence. Plot shows the biophysical preference for disease causing and neutral phospho variants for central site (P) and its flanking sites (P-5 to P5). KS statistics are shown only for the data points at central site ( position ‘P’).

**A)** Distribution of Backbone dynamic values. Values > 0.8: rigid, 0.69-0.8: context dependent, <0.69: flexible. **B)** Distribution of disorder propensity values. Values above 0.5indicate that this is likely a disordered residue. **C)** Sidechain dynamics: Note at ‘P’ its just S/T/Y residues and the flanking residues can be any amino acid. Higher values mean more likely rigid. These values are highly dependent on the amino acid type (i.e. a Trp will be rigid, an Asp flexible). **D)** Early folding propensity. Values above 0.169 indicate residues that are likely to start the protein folding process, based on only local interactions with other amino acids.

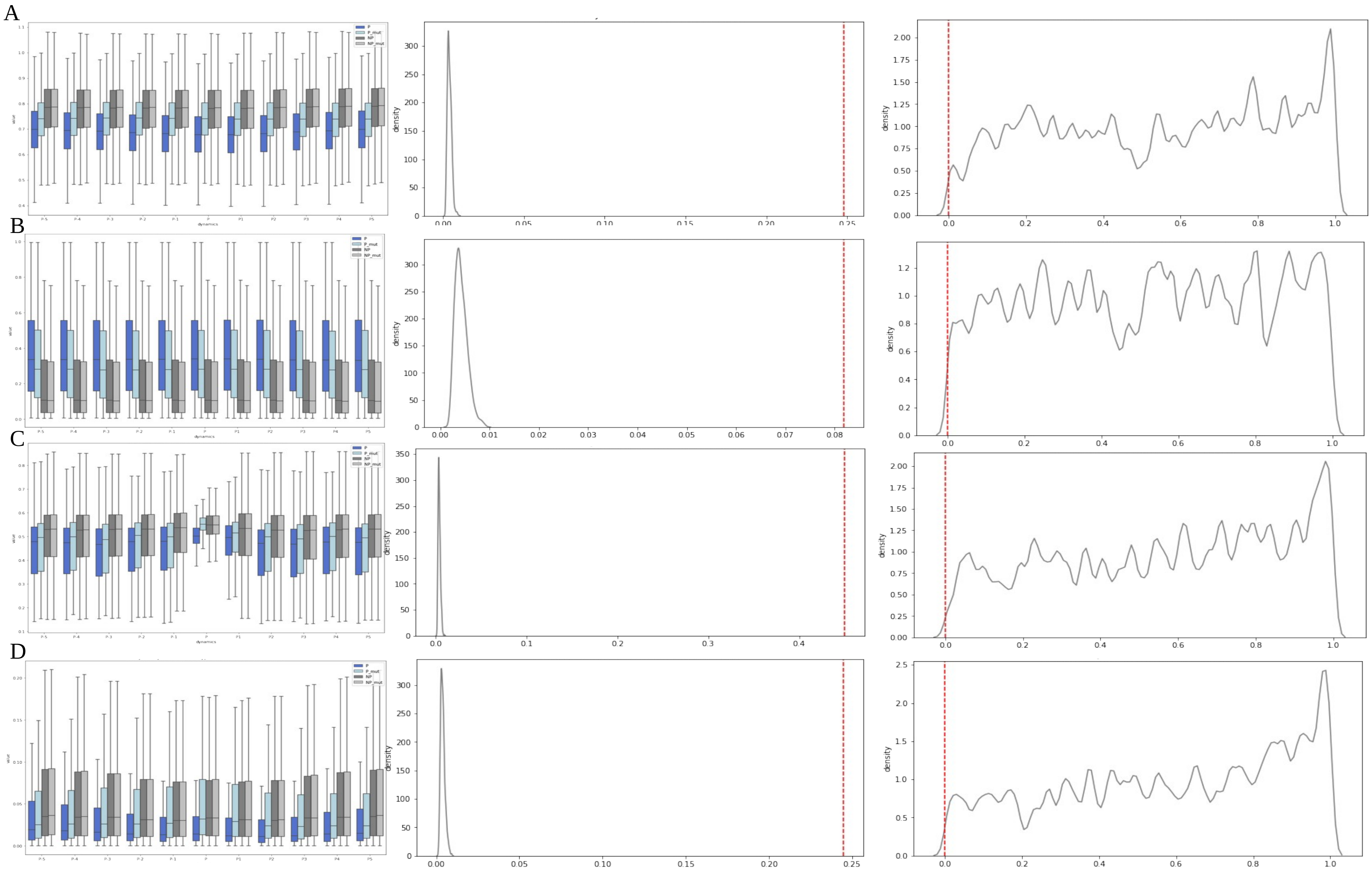

**Figure S7: Biophysical properties of P-sites and mutated S>A,T>V,Y>F at P-sites and non-P-sites**

**Left panel** -Distribution of dynamic values of P-sites (blue, N=79994), mutated phospho STY>AVF and non-P-sites (dark and light grey, N= 1116380). **Middle panel** shows the KS-Dstat distribution of 1000 random permutation test by shuffling P-site and mutated STY sites (red line is the actual observed value between two group) and **right panel** shows the P-value distribution of random permutation test (red line is the actual observed value) Predictions are done on primary amino acid sequence. Plot shows the biophysical preference for P-sites (blue), mutated P-sites (light blue), non P-sites in wild type sequence (dark grey) and non-P-sites in mutated sequence (light grey) for central site (P) and its flanking sites (P-5 to P5). KS statistics are shown for the P-sites (blue) and mutated P-site (light blue) groups only (position ‘P’).

**A)** Distribution of Backbone dynamic values. Values > 0.8: rigid, 0.69-0.8: context dependent, <0.69: flexible. **B)** Distribution of disorder propensity values. Values above 0.5indicate that this is likely a disordered residue. **C)** Sidechain dynamics: Note at ‘P’ its just S/T/Y residues and the flanking residues can be any amino acid. Higher values mean more likely rigid. These values are highly dependent on the amino acid type (i.e. a Trp will be rigid, an Asp flexible). **D)** Early folding propensity. Values above 0.169 indicate residues that are likely to start the protein folding process, based on only local interactions with other amino acids.

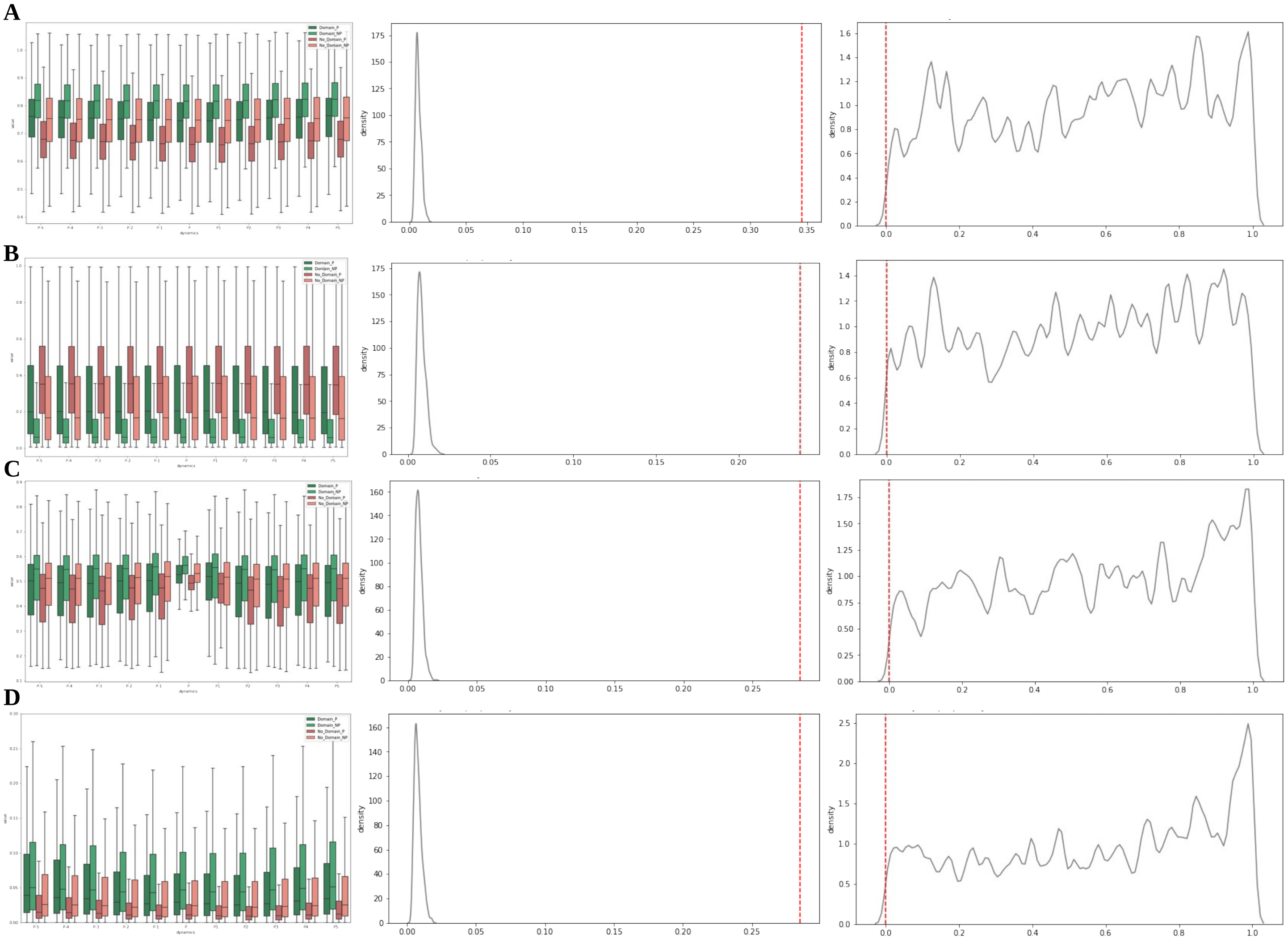

**Figure S8: Biophysical properties of P-sites and non-P-sites in domain region and those not in domain regions**
**Sites in domain region are colored in green shades and sites not in domain regions are colored in red shades. P-sites are indicated by dark colors (green and Indian red)**
**Left panel** -Distribution of dynamic values of P-sites in domain (dark green, N=20639) and not in domain (Indian red, N= 30991), non P-sites in domain (light green, N=330916) and non-P-site not in domain (salmon, N=278876). **Middle panel** shows the KS-Dstat distribution of 1000 random permutation test by sampling (every permutation) equal size of P-site not in domain and shuffling (red line is the actual observed value between two group) and **right panel** shows the P-value distribution of random permutation test (red line is the actual observed value) Predictions are done on primary amino acid sequence. Plot shows the biophysical preference for the sites to be phosphorylated in domains (P-sites) and not in domain (P-sites not in domain) for central site (P) and its flanking regions (P-5 to P5). KS statistics are shown for the central P-sites with domain (dark green) and P-sites in non domain regions (Indian red) at position ‘P’

**A)** Distribution of Backbone dynamic values. Values > 0.8: rigid, 0.69-0.8: context dependent, <0.69: flexible. **B)** Distribution of disorder propensity values. Values above 0.5indicate that this is likely a disordered residue. **C)** Sidechain dynamics: Note at ‘P’ its just S/T/Y residues and the flanking residues can be any amino acid. Higher values mean more likely rigid. These values are highly dependent on the amino acid type (i.e. a Trp will be rigid, an Asp flexible). **D)** Early folding propensity. Values above 0.169 indicate residues that are likely to start the protein folding process, based on only local interactions with other amino acids.

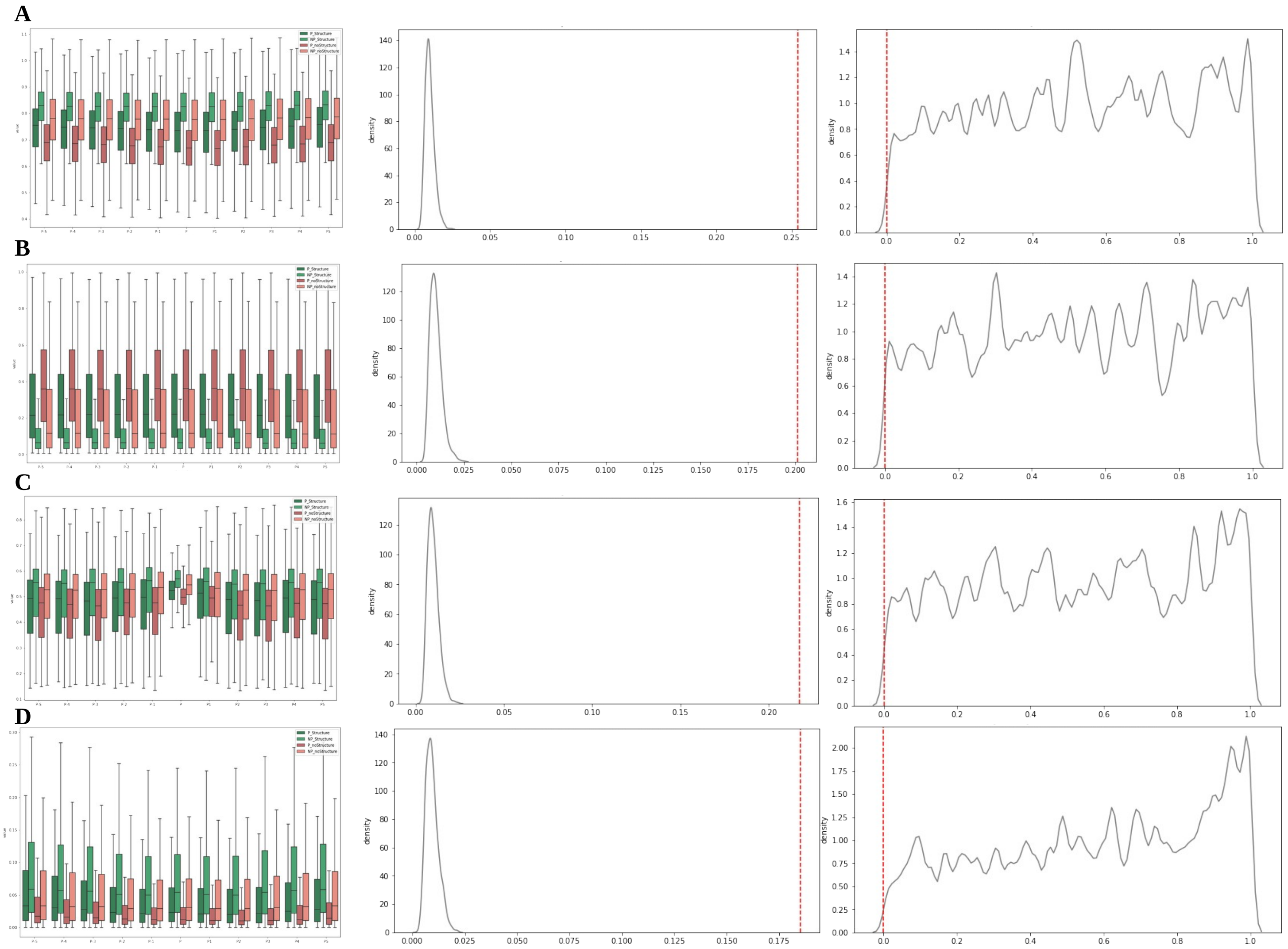


**Figure S9: Biophysical properties of P-sites and non-P-sites with and without structural annotations**
**Sites with structural annotations are colored in green shades and sites with no structural annotation are colored in red shades. P-sites are indicated by dark colors (green and Indian red)**
**Left panel** -Distribution of dynamic values of P-sites in domain (dark green, N=13969) and not in domain (Indian red, N= 66025), non P-sites in domain (light green, N=164430) and non-P-site not in domain (salmon, N=1086461). **Middle panel** shows the KS-Dstat distribution of 1000 random permutation test by sampling (every permutation) equal size of P-site with no structures and shuffling (red line is the actual observed value between two group) and **right panel** shows the P-value distribution of random permutation test (red line is the actual observed value) Predictions are done on primary amino acid sequence. Plot shows the biophysical preference for P-sites non-P-sites with and with no structural annotation for central site (P) and its flanking regions (P-5 to P5). KS statistics are shown for the central P-sites with structural annotation (dark green) and P-sites with no structural annotation (Indian red) at at position 'P'

**A)** Distribution of Backbone dynamic values. Values > 0.8: rigid, 0.69-0.8: context dependent, <0.69: flexible. **B)** Distribution of disorder propensity values. Values above 0.5indicate that this is likely a disordered residue. **C)** Sidechain dynamics: Note at 'P' its just S/T/Y residues and the flanking residues can be any amino acid. Higher values mean more likely rigid. These values are highly dependent on the amino acid type (i.e. a Trp will be rigid, an Asp flexible). **D)** Early folding propensity. Values above 0.169 indicate residues that are likely to start the protein folding process, based on only local interactions with other amino acids.

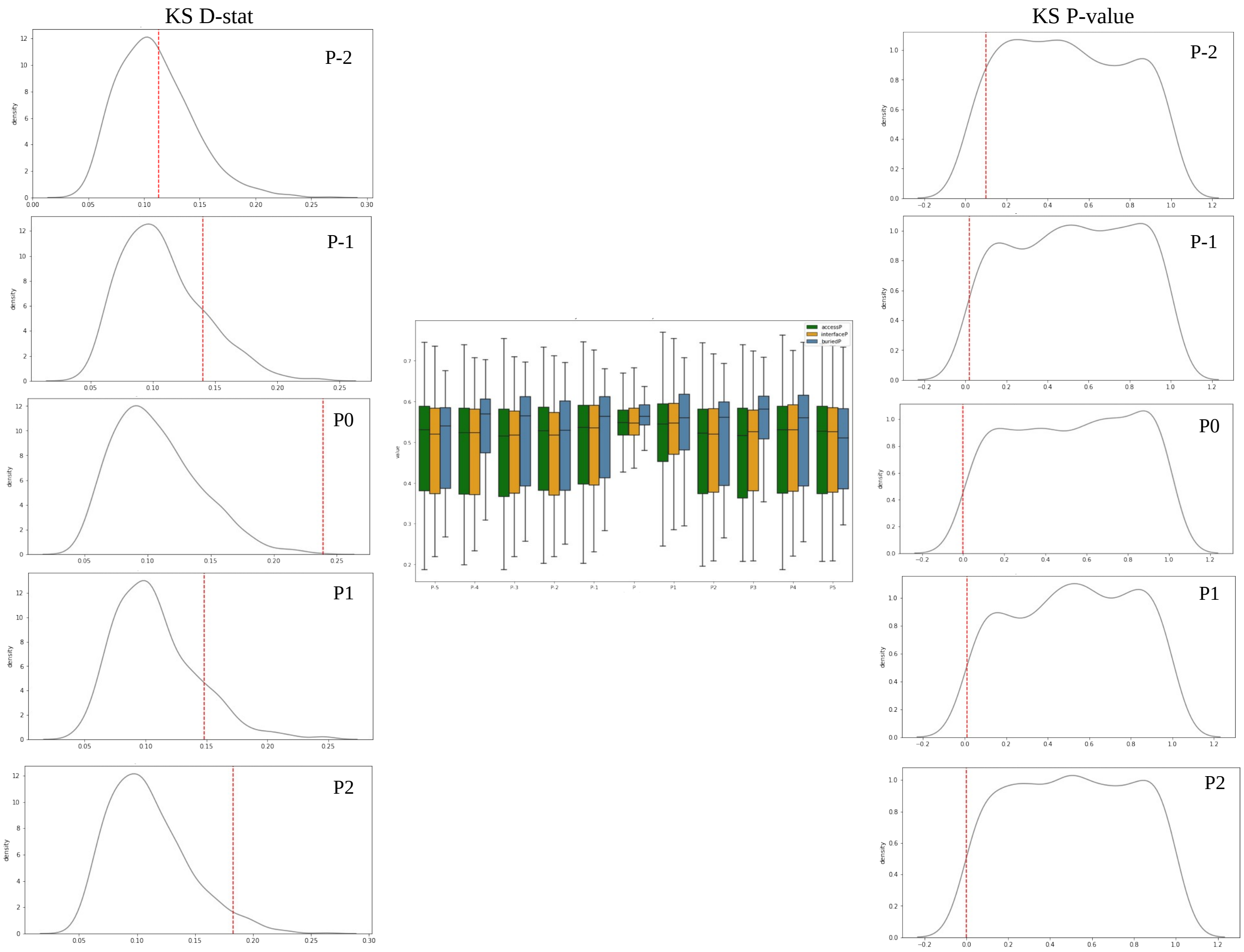


**Figure S10: Side chain dynamics of central P-sites and flanking positions**

**Middle panel** -Distribution of dynamic values of P-sites in solvent exposed structural regions (green, N=5406), interface regions (orange, N=990) and in buried/in-accessible regions (blue, N=116). **Left panel** shows the KS-Dstat distribution of 1000 random permutation test by sampling (every permutation) P-sites in accessible region to match the sample size in buried regions (116), maintained equal size for residues S:74, T:31, Y:11 and shuffling. (red line is the actual observed value between two group) .and **right panel** shows the P-value distribution of random permutation test (red line is the actual observed value) Predictions are done on PDB sequence. Plot shows if there is any biophysical preference for some P-sites in accessible, interface and buried regions of the protein structures. KS statistics are shown for the central sites P and flanking positions 'P-2','P-1','P1','P2'. Side chain dynamics of accessible and buried residues in N terminal region (P-1, P-2) are very similar and a slight yet significant similarity of side chains observed at C terminal flanking sites (P1, P2).

**Side chain dynamics:** Note at 'P' its just S/T/Y residues and the flanking residues can be any amino acid. Higher values mean more likely rigid. These values are highly dependent on the amino acid type (i.e. a Trp will be rigid, an Asp flexible).

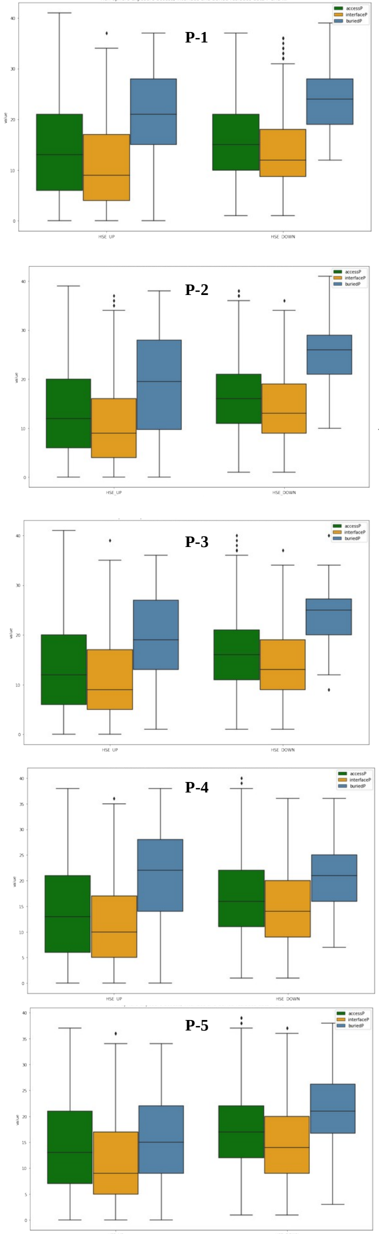


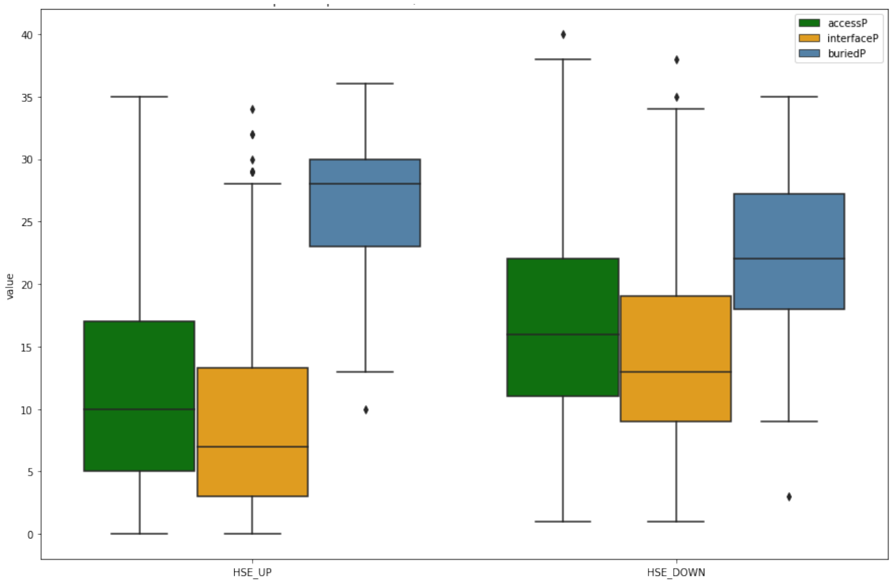


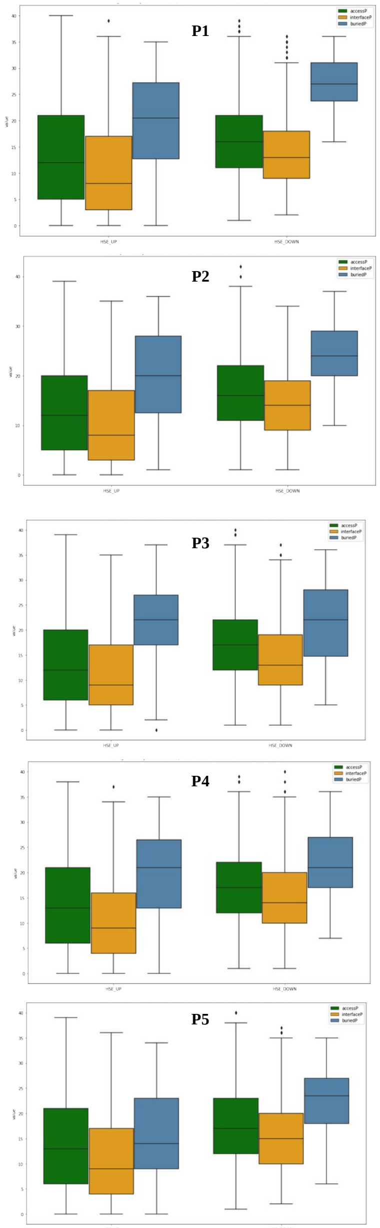


**Figure S11: Half sphere exposure (HSE) of P-sites in solvent accessible, interface and buried regions**

**Middle panel:** Number of Cβ contacts in the direction of side chain (HSE_UP) and in opposite direction (HSE_DOWN) for the P-sites which are seen in solvent accessible (green), interface (orange) and buried (blue) regions of the protein structures. Buried residues shows more residue contacts in side chain directions and in the direction of backbone (HSE_UP).

**Left Panel:** Number of Cβ contacts for the N-terminal flanking residues (P-1 to P-5). Slight decrease in tye median values (black bar in box plots) of HSE_UP contacts and a slight increase in HSE_down median contacts (black bar in box plots) and at P-5 the HSE_UP for accessible and buried residues are almost similar (p-value 0.06)

**Right Panels:** Number of Cβ contacts for the C-terminal flanking residues (P1 to P5). Like N terminal the C terminal sites shows a slight decrease in HSE_UP median contacts (black bar in box plots) and a slight increase in HSE_down median contacts (black bar in box plots) and at P5 the HSE_UP for accessible and buried residues are almost similar (p-value 0.052)

Though the residue contacts of accessible and interface residues are consistent, the contacts of buried residues increases with decrease in flanking distance.

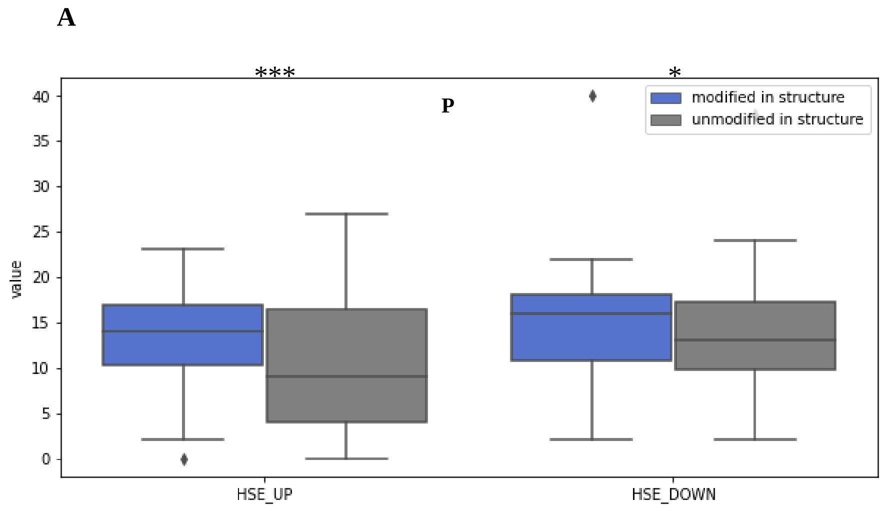


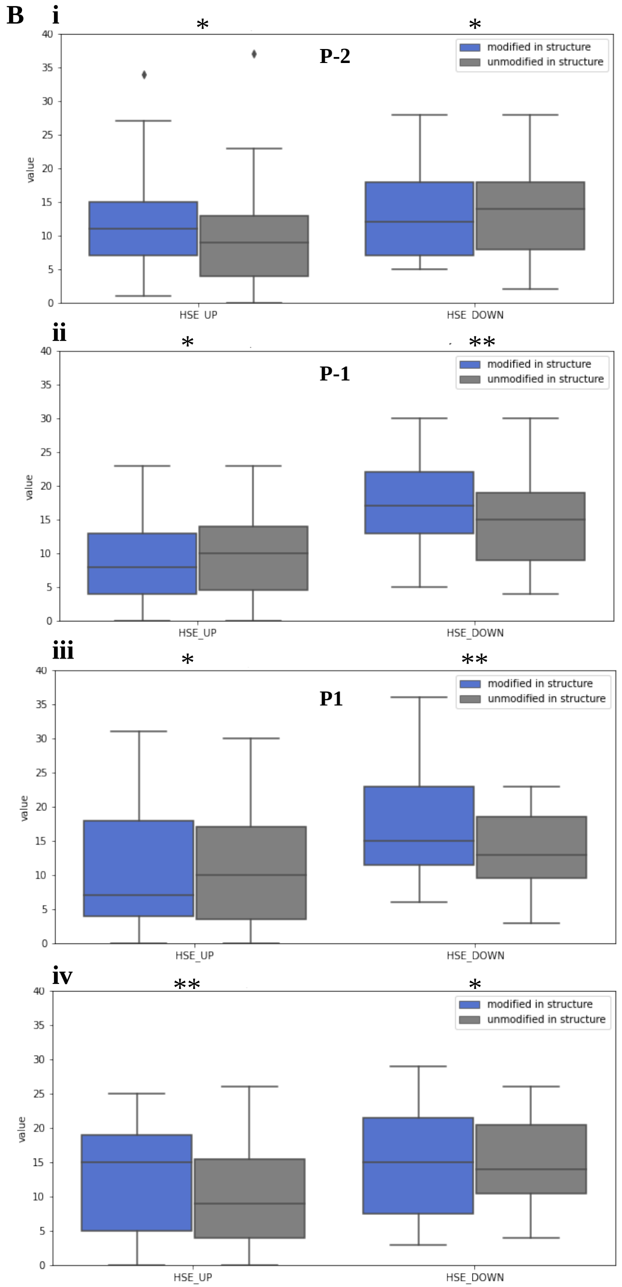


**Figure S12: Half sphere exposure (HSE) of sites with and without phosphorylation**

**A)** Number of Cβ contacts in the direction of side chain (HSE_UP) and in opposite direction (HSE_DOWN) for the sites which are seen as phopshorylated in structure (blue) and same site seen as un-phopshorylated in a different structure (grey) from same proteins. B**)** Number of Cβ contacts in structure for the flanking residues in sequence P-2 (**i**), P-1 (**ii**), P1**(iii)** and P2 **(iv)** color represents the same as in A.

***** p-value <0.05, ** p-value <0.5, * p-value >0.5**
